# Psilocybin modulates social behaviour in male and female mice in a time-dependent manner

**DOI:** 10.64898/2025.12.18.695064

**Authors:** Sheida Shadani, Kaspar McCoy, Lina Ong, Erika Greaves, Kyna Conn, Zane B. Andrews, Claire J. Foldi

## Abstract

With the resurgence of psychedelic research and the growing interest in their therapeutic potential, there is an urgent need to understand how these compounds act across biological sexes. Despite widespread interest in their use for conditions marked by social impairments, including depression, anxiety, and anorexia nervosa, the influence of sex as a biological variable (SABV) on the prosocial effects of psychedelics remains poorly understood. Indeed, enhanced connectedness, sociability and empathy are common outcomes of psychedelic use and these have shaped human social structures for millennia. Here, we investigated the sex-specific effects of a single dose of psilocybin (1.5 mg/kg) in C57BL/6J mice on various aspects of social behaviours. We show an intriguing connection between huddling behaviour and body temperature acutely elicited by psilocybin that was restricted to females. We also observe temporally distinct patterns of social behaviour alterations in female mice, whereby enhanced preference for social novelty was observed after acute effects subsided (4 h post-administration), which was maintained for ∼24 h. Longer-term, the impact of psilocybin was reversed and promoted preference for familiar over novel conspecifics when assessed 7d post-administration, which was associated with prolonged nucleus accumbens dopamine signalling during familiar sniffing. In males, psilocybin reduced stress-related behaviours at 24 h and increased preference for familiar conspecifics, along with blunted novelty-evoked dopamine responses at both 24 h and 7 days post-treatment. Both 5-HT1A and 5-HT2A receptors were involved in modulating these behaviours, though in sex-specific ways. These findings highlight that the prosocial effects of psychedelics are not universal and emphasize the importance of sex-informed approaches in both preclinical research and clinical application.

**Graphical Abstract:** 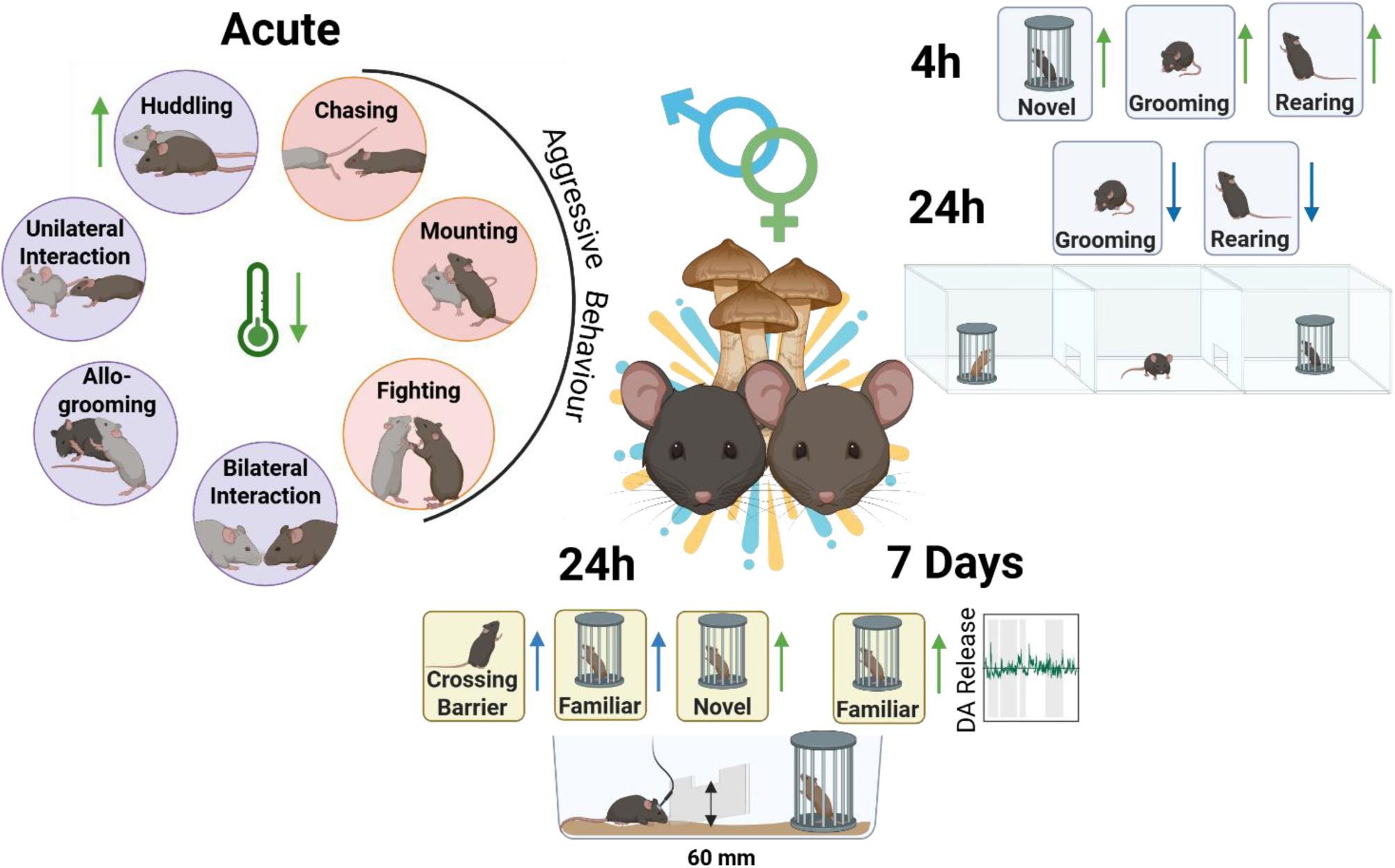

## Introduction

Social behaviour is a fundamental to human cognition, shaping societal networks through cooperation, affiliation, and trust [1]. The ability to form and maintain social bonds is crucial for psychological well-being, and social deficits characterise numerous psychiatric and neurological disorders, including autism spectrum disorder, schizophrenia, depression, and anorexia nervosa [2, 3]. Emerging evidence suggests that psychedelics, particularly psilocybin, may enhance social cognition and promote prosocial behaviour by modulating neural circuits involved in social reward and emotional processing [4, 5].

The resurgence of psychedelic research has revealed therapeutic potential for major depressive disorder, generalised anxiety disorder, obsessive-compulsive disorder, post-traumatic stress disorder (PTSD) and anorexia nervosa [6–14]. However, despite the central role of social behaviour in mental health, the impact of psychedelics on social cognition and motivation remains relatively underexplored. Current research has focused on primary symptom reduction in clinical populations, overlooking the potential for psychedelics to enhance social functioning that may be critical for long-term therapeutic efficacy. Expanding research into the social effects of psychedelics could offer novel mechanistic insights and refine clinical applications in disorders characterized by social impairments.

Emerging evidence suggests that psychedelics such as lysergic acid diethylamide (LSD) and psilocybin can enhance social connectivity in humans by modulating affective processing and social cognition [15]. Psilocybin has been shown to improve recognition of negative emotional stimuli [16], reduce feelings of social exclusion [17], and enhance empathy and prosocial behaviours [18], though these effects are not consistently observed across all studies [19]. These effects have been observed in both healthy individuals and patients with major depressive disorder [16, 17, 20, 21], supporting a role for psilocybin in fostering interpersonal connectedness [22]. Furthermore, single doses of psilocybin produce long-lasting improvements in social functioning and mood [23–25], suggesting persistent neuroplastic adaptations that enhance social engagement [26, 27]. Healthy volunteers also report elevated positive mood and prosocial behaviours under acute psilocybin effects [4, 28, 29], reinforcing its potential as a modulator of affective and social processing.

Preclinical studies investigating the influence of psychedelics on social behaviour in rodents show mixed outcomes. Psilocybin elicits both prosocial and antidepressant-like effects in male rats [30], while LSD and psilocybin both increase social interaction, preference for social novelty [31–34] and social reward learning [35] in male mice. However, a recent multi-institutional study failed to replicate effects of psilocybin on social reward learning or preference for novel social stimuli in male and female mice [36]. Notably, neither study reported or discussed potential sex-specific differences in sociability. Underscoring the need for standardised protocols, reproducibility efforts, and inclusion and separate analysis of both sexes to evaluate sex-dependent effects.

While interest in the behavioural effects of psychedelics has increased, underlying neurobiological mechanisms remain inadequately understood. The medial prefrontal cortex (mPFC), crucial for social decision-making and regulation of prosocial behaviours [37–39], is the most commonly reported site of psychedelic effects in human imaging and animal studies. However, psilocybin also modulates activity within the anterior cingulate cortex (ACC), involved in social pain processing [17], helping behaviour and empathy [40, 41] and social decision making [42], as well as the amygdala, which regulates social anxiety and threat perception [43]. These regional effects are linked to interactions with the mesolimbic dopamine system and the oxytocin producing pathways, both of which are known to regulate pair bonding [44]. For instance, MDMA reopens social reward learning windows via oxytocin receptor activation in the nucleus accumbens [45], while LSD and psilocybin modulate connectivity between the default mode network (DMN) and social processing hubs, potentially reducing egocentric bias and increasing social connectedness [32]. Rodent studies further implicate the paraventricular nucleus of the hypothalamus (PVN), a key oxytocin producing site, as a target for psychedelic-induced social facilitation [46]. Collectively, these findings suggest that psychedelics reshape social behaviour by engaging cortical and subcortical circuits that regulate social motivation, reward, and emotional salience.

Despite promising findings, the majority of psychedelic research in humans and animal models has disproportionately (or often times exclusively) focused on male participants, hindering generalizability given well-documented sex differences in neurobiology, psychiatric disease prevalence, and pharmacological responses [47, 48]. For instance, selective serotonin reuptake inhibitors (SSRIs), which target the serotonergic system key to psychedelic actions, exhibit sex-dependent efficacy and side-effect profiles [49]. Moreover, conditions such as depression, anxiety, PTSD, and eating disorders, many being potential targets for psychedelic therapy, are significantly more prevalent in females [50–52]. To address this critical gap and align with the National Institutes of Health mandate to integrate sex as a biological variable (SABV) in experimental design [53], the present study investigated the effects of psilocybin on multiple aspects of social behaviour in both male and female mice and across different time points (from acute to post-acute administration). In addition, we tested the hypothesis that psilocybin binding to specific serotonin (5-HT) receptor subtypes is necessary for eliciting prosocial behaviour and paired social behaviour tasks with *in vivo* fiber photometry to investigate the neuronal substrates underlying these effects.

## Methods

### Animals and housing

Male (*n*=182) and female (*n*= 145) C57Bl/6 mice (10 weeks old) were obtained from the Monash Animal Research Platform and pair-housed under reverse light cycle conditions (lights off at 0800h). Mice acclimated for 7 days before testing, which occurred between 1000h and 1700h. Each mouse underwent only one experimental condition. All experimental procedures were approved by the Monash Animal Research Platform Ethics Committee (ERM 30852, ERM 40619).

### Drug administration

Psilocybin (PSI; 1.5mg/kg) was administered intraperitoneally in saline. In receptor antagonist experiments, MDL100907 (5-HT2A antagonist; 0.1mg/kg) or WAT100635 (5-HT1A antagonist; 0.5mg/kg) were administered 30 mins prior to PSI or saline (SAL) control. Doses were selected based on previous studies [54]. See **Supplementary Methods** for drug preparation and supplier details.

### Behavioural testing

#### Home-cage social behaviour

Following PSI or SAL injection, pair-housed mice were video-recorded for 1 h in their home cage. Social behaviours including bilateral interaction, huddling, allogrooming, mounting, and chasing were manually scored using Ethovision XT software.

#### 3-Chamber test

Mice were tested 4 h, 24 h, or 7 d post-injection using the standard 3-chamber paradigm. After 10 min habituation, mice explored a novel mouse versus empty cage (social preference trial), then a second novel mouse versus the now-familiar mouse (social novelty trial). Time in chambers, interaction frequency, and locomotor activity were quantified. Social preference was calculated as (T_s_ - T_ns_) / (T_s_ + T_ns_), where T_s_ is time spent with the novel mouse, and T_ns_ is time spent with the empty cage or familiar mouse. Sociability Z-scores were calculated by normalizing individual scores to control group means: Z = (x - μ)/σ, based on chamber duration and entry frequency [55].

#### Barrier climbing test

Mice were assessed for social motivation using an adapted maternal motivation paradigm [56–58]. Following 10 min habituation to a 60 mm transparent barrier, mice explored a novel mouse (5 min) then a cage mate (5 min), each confined in a wire cage on the opposite side of the barrier. Latency to first climb, total climbs, and latency to sniff were recorded.

#### Acute physiological monitoring

Core body temperature and locomotor activity were continuously monitored using subcutaneously implanted RFID microchips (UCT-2112; Unified Information Devices) in the home cage for 1 h before and 2 h after injection.

### Fiber photometry

Mice received unilateral injections of hSyn-GRAB_DA2m_ (Addgene #140553) and optical fiber implantation (AP:+1.4, ML: ±0.75, DV:-4.2 mm). Following 5 weeks recovery, dopamine dynamics were recorded during the barrier climbing task using 470 nm (signal) and 410 nm (isosbestic control) excitation within the RWD R821 system. See **Supplementary Methods** for extended surgical and viral injection procedure details.

### Statistical analyses

Data were analysed using GraphPad Prism 9.5.1 with significance set at *p*< 0.05. Analyses included unpaired t-tests, one-way and two-way ANOVA with Sidak’s post hoc tests, and mixed effects models as appropriate. Complete statistical details are provided in the **Supplementary Materials.**

## Results

### Psilocybin-induced social thermoregulation in female mice is 5-HTR dependent

To evaluate the acute effects of psilocybin on home-cage social behaviour, we examined huddling in pair-housed female mice. Psilocybin significantly increased huddling within 60 min post-administration (*p* = 0.0391; **Figure 1A**), with peak effects at 25 min (*p* = 0.0133; **Figure 1B**). This coincided with significant reductions in core body temperature beginning at 5 min (*p* = 0.0277) and persisting until 70 min (*p* = 0.0073; **Figure 1C**). 5-HT1AR antagonism abolished huddling regardless of subsequent treatment (saline: *p* = 0.0062; psilocybin: *p* < 0.0001), while 5-HT2AR antagonism selectively reduced huddling in psilocybin-treated mice (*p* = 0.0029; **Figure 1D**). In saline-treated mice, 5-HTR antagonism did not affect huddling (**Figure 1E**); however, in psilocybin-treated mice, 5-HT1AR antagonism abolished huddling at its 30 min peak (*p* = 0.0158; **Figure 1F**). Both 5-HT1AR and 5-HT2AR antagonism reduced core body temperature in saline-treated controls, more pronounced and longer lasting for 5-HT1AR (*p* = 0.0244), with only two timepoints significantly altered following 5-HT2AR antagonism (45 min: *p* = 0.0258 and 65 min: *p* = 0.0099; **Figure 1G**). In psilocybin-treated mice, 5-HT1AR antagonism prolonged temperature reduction, whereas 5-HT2AR antagonism acutely increased temperature (*p* = 0.0398), remaining elevated above psilocybin-only levels between 60-90 min (85 min: *p* = 0.0429; **Figure 1H**).

**Figure 1.**
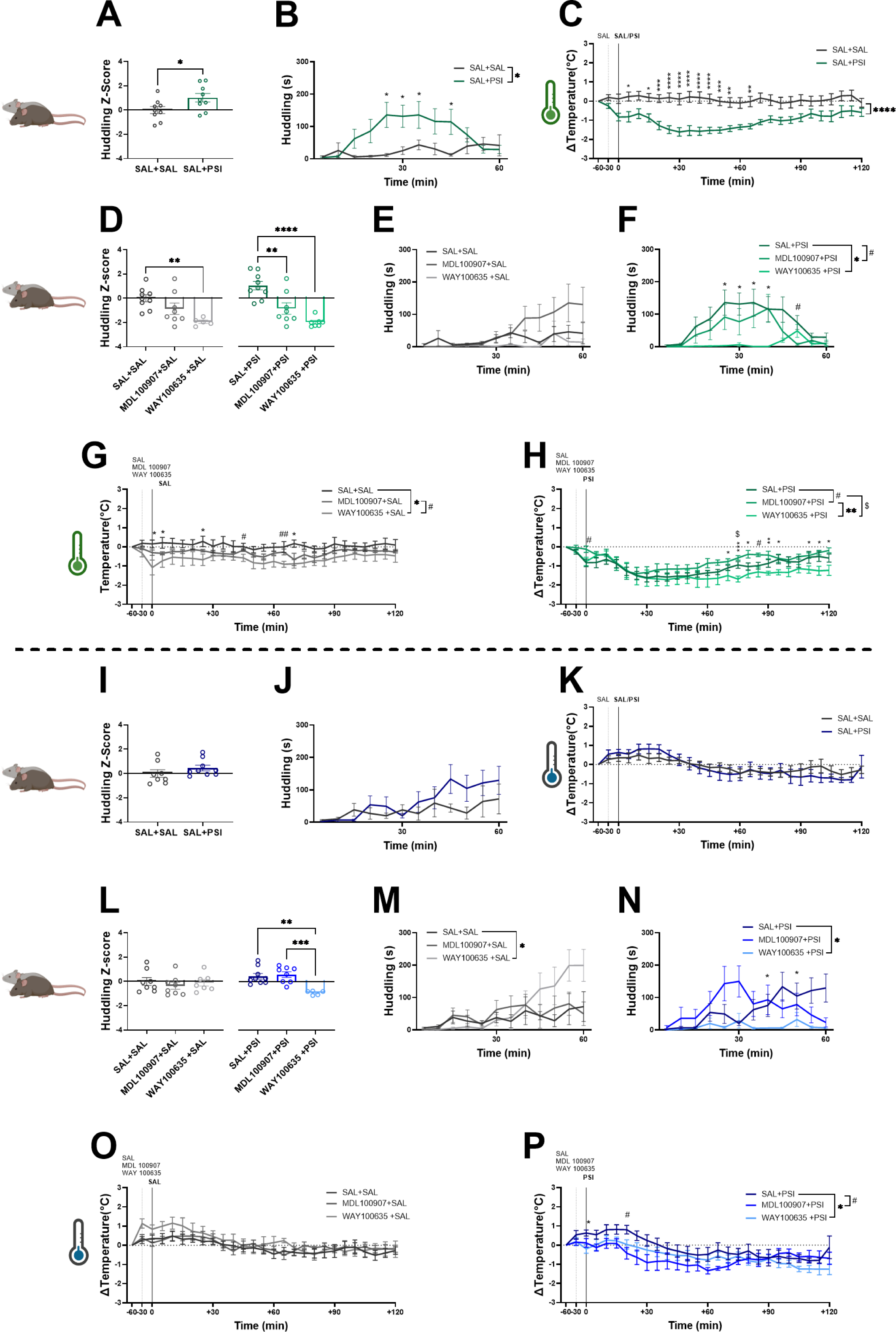
Psilocybin increases huddling in females but not males, while 5-HT1A and 5-HT2A receptor antagonism modulate huddling and temperature in a sex-specific manner. **(A)** Psilocybin significantly increased huddling behaviour compared to the controls (*p* = 0.0391) with **(B)** the huddling behaviour peaking at 25 min (*p* = 0.0133), 30 min (*p* = 0.0155) and 45 min (*p* = 0.0311) post-administration. **(C)** Psilocybin significantly reduced core body temperature in a time-dependent manner between 5-70 minutes post-administration (*F* (1,160) = 16.1, *p* = 0.001). **(D)** Among SAL-treated mice, 5-HT1AR antagonism reduced huddling behaviour (*p* = 0.0062) and in PSI-treated mice, both 5-HT1AR (*p* < 0.0001) and 5-HT2AR (*p* = 0.0029) antagonists reduced huddling behaviour. **(E)** 5-HTR antagonism did not significantly affect huddling behaviour over the 60 minutes observation period in SAL-treated mice however, **(F)** in PSI-treated mice, 5-HT1AR antagonism reduced huddling behaviour at 25 min (*p* = 0.0228), 30 min (*p* = 0.0158), 35 min (*p* = 0.0275) and 40 min (*p* = 0.0428) while 5-HT2AR reduced huddling behaviour only at 50 min (*p* = 0.0419) compared to PSI-treated group. **(G)** In SAL-treated mice, 5-HT1AR antagonism lowered core body temperature at 0 min (*p* = 0.0206), 5 min (*p* = 0.0398), 25 min (*p* = 0.0257), 65 min (*p* = 0.027) and 70 min (*p* = 0.0112) compared to SAL-treated group and it produced a stronger temperature reduction than 5-HT2AR antagonist at 45 min (*p* = 0.0258) and 65 min (*p* = 0.0099). **(H)** In PSI-treated mice, 5-HT1AR reduced the core body temperature at 75 min (*p* = 0.021), while 5-HT2AR increased temperature at 0 min (*p* = 0.0398) and 85 min (*p* = 0.0429) compared to PSI-treated group. Additionally, 5-HT1AR antagonism led to significantly lower body temperature than 5-HT2AR antagonism between 75-120 minutes (Time x Treatment interaction: *F* (2,22) = 2.868, *p* = 0.0152). **(I)** In male mice, total huddling behaviour was not significantly affected by psilocybin and **(J)** no time-dependent differences in huddling behaviour were observed over 60 minutes. **(K)** Core body temperature was not significantly altered by psilocybin. **(L)** In SAL-treated male mice, 5-HTR antagonism did not significantly affect the huddling behaviour. However, in PSI-treated mice, 5-HT1AR antagonism significantly reduced huddling behaviour compared to PSI only group (*p* = 0.0018) and compared to the 5-HT2AR antagonism (*p* = 0.0009). **(M)** No significant differences in huddling behaviour were observed over 60 minutes among saline-treated groups. **(N)** In PSI-treated mice, 5-HT1AR antagonism reduced huddling behaviour at 45 min (*p* = 0.0496) and 55 min (*p* = 0.0489). **(O)** Core body temperature did not differ significantly among saline-treated groups over 60 minutes. **(P)** In PSI-treated mice, 5-HT1AR reduced core body temperature at 0 min (*p* = 0.0289) and 5-HT2AR antagonism also lowered core body temperature at 20 min (*p* = 0.0165). Female mice (A-H), male mice (I-P), psilocybin (PSI), saline (SAL), 5-HT2AR antagonist (MDL100907), 5-HT1AR antagonist (WAY100635). Data are presented as mean ± SEM. Statistical analyses were performed using unpaired t-test, mixed-effects models, one-way ANOVA or two-way ANOVA with Šidák post hoc tests. \**p*< 0.05, ^#^*p*< 0.05, ^$^*p*< 0.05, \*\**p*< 0.01, ^##^*p*< 0.01, \*\*\**p*< 0.001, \*\*\*\**p*< 0.0001.

Unilateral interaction was significantly reduced by 5-HT2AR antagonism, but not by 5-HT1AR antagonism, in pair-housed females regardless of subsequent treatment (saline: *p* = 0.0339; psilocybin: *p* = 0.0328). Neither psilocybin nor 5-HTR antagonism affected aggressive behaviour in female mice (**Supplementary Data 1**). Within the groups pre-treated with 5-HTR antagonists, psilocybin reduced core body temperature in mice treated with 5-HT2AR antagonists at 45 min (*p* = 0.0297), and in 5-HT1AR-antagonised mice at 75 min (*p* = 0.0446) and 90 min (*p* = 0.0478; **Supplementary Data 2**).

### Huddling behaviour in psilocybin-treated male mice is modulated by 5-HTR antagonism

Psilocybin alone did not alter huddling behaviour or core body temperature in male mice (**Figure 1I-K**), nor did 5-HTR antagonism in saline-treated males (**Figure 1L-M**, **Figure 1O**). However, 5-HT1AR antagonism significantly reduced huddling in psilocybin-treated males (*p* = 0.0018; **Figure 1L**), consistently observed across the 60 min monitoring period (*p* = 0.0399; **Figure 1N**). Combined with psilocybin, both 5-HTR antagonists reduced core body temperature compared to psilocybin alone (*p* = 0.0172; **Figure 1P**). Neither psilocybin nor 5-HTR antagonism affected unilateral interactions in males. However, both 5-HT1AR and 5-HT2AR antagonism significantly reduced aggressive behaviour (primarily mounting, followed by chasing, pinning and fighting), effects absent when 5-HTR antagonists were combined with psilocybin (**Supplementary Data 1**). Comparing mice receiving psilocybin or saline with 5-HTR antagonists revealed opposing effects: For 5-HT2AR antagonist recipients, co-administration of psilocybin increased huddling (*p* = 0.0194), whereas for 5-HT1AR antagonist recipients, co-administration of psilocybin reduced huddling (*p* = 0.0327). Both psilocybin-treated groups receiving 5-HTR antagonists exhibited reduced core body temperature compared to their saline-treated counterparts (5-HT1AR: *p* = 0.0193; 5-HT2AR: *p* = 0.0096; **Supplementary Data 3**).

### Psilocybin modulates novelty-seeking behaviour in female mice through 5-HT1AR and 5-HT2AR mechanisms

Using the 3-chamber test to assess how psilocybin impacts social preference and novelty-seeking behaviour in mice (**Figure 2A**), we observed effects in female mice at 4 h post-administration unrelated to changes in locomotor activity (**Figure 2B**). Psilocybin enhanced novelty-seeking with a trend toward increased sociability compared to controls (*p* = 0.097; **Figure 2C**) and a significant preference for the novel mouse (*p* = 0.0494; **Figure 2D**). Psilocybin significantly increased grooming (*p* = 0.0443; **Figure 2E**), particularly during the choice phase (*p* = 0.0514; **Supplementary Data 4**). Rearing behaviour was not significantly affected (**Figure 2F**). Both 5-HT1AR and 5-HT2AR antagonism attenuated psilocybin-associated sociability (5-HT1AR: *p* = 0.023; 5-HT2AR: *p* = 0.0252), but neither altered sociability in saline-treated controls (**Figure 2G**). This pattern was similarly reflected in direct sociability measures (5-HT1AR: *p* = 0.0243; 5-HT2AR: *p* = 0.0007; **Figure 2H**). Notably, 5-HT2AR antagonism blocked psilocybin-induced sociability without affecting locomotor activity, supporting a specific role for 5-HT2AR at 4 h. In contrast, 5-HT1AR antagonism reduced locomotion at this timepoint (**Supplementary Data 4**), suggesting suppressed sociability reflects reduced general activity rather than a direct social effects. Therefore, the role of 5-HT1AR in mediating psilocybin’s prosocial effects requires cautious interpretation. In males, neither locomotion nor sociability measures were affected by psilocybin at 4 h (**Supplementary Data 5**).

**Figure 2.**
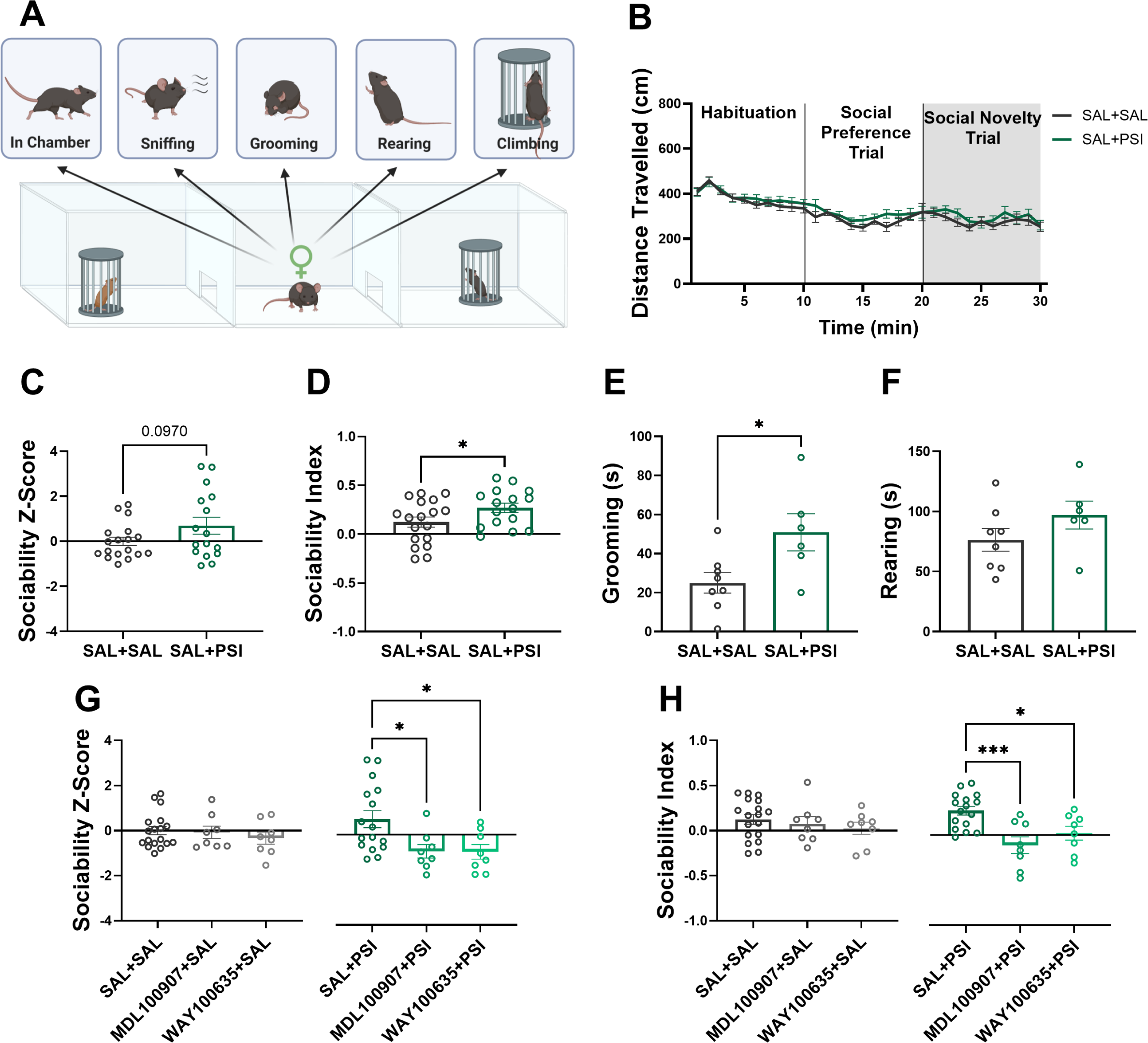
Psilocybin enhances sociability in female mice 4 hours after administration, and these effects are blocked by 5-HT receptor antagonism. **(A)** Schematic of the social novelty trial and the behaviours scored. **(B)** Psilocybin did not alter locomotor activity of mice across the duration of the experiment. **(C)** No significant differences were observed in the total sociability Z-score between the groups (*p* = 0.097). **(D)** PSI-treated mice displayed significantly higher sociability compared to controls (*p* = 0.0494). **(E)** Psilocybin increased grooming behaviour (*p* = 0.0443) but **(F)** did not affect the rearing behaviour in these mice. **(G)** In SAL-treated mice, 5-HTR antagonism did not affect sociability. In contrast, in PSI-treated mice, both 5-HT1AR (*p* = 0.023) and 5-HT2AR (*p* = 0.0252) antagonism significantly reduced sociability Z-score. **(H)** Similarly, 5-HTR antagonism did not affect sociability in SAL-treated mice but reduced sociability in PSI-treated mice with both 5-HT1AR (*p* = 0.0243) and 5-HT2AR (*p* = 0.0007) antagonism. Psilocybin (PSI), saline (SAL), 5-HT2AR antagonist (MDL100907), 5-HT1AR antagonist (WAY100635). Data are presented as mean ± SEM. Statistical analyses were performed using unpaired t-test, one-way ANOVA or two-way ANOVA with Šidák post hoc tests. *p< 0.05, ***p< 0.001.

### Psilocybin attenuates rearing and grooming in male mice at 24 hours post-administration, but does not alter sociability

Given the possibility that psilocybin’s time course differs between sexes, we conducted the 3-chamber test in male mice at 24h (**Figure 3A**). Psilocybin did not affect locomotor activity (**Figure 3B**) or sociability measures (**Figure 3C, D**) but significantly reduced grooming (*p* = 0.0159; **Figure 3E**) and rearing (*p* = 0.0033; **Figure 3F**). Unlike saline-treated females, 5-HT2AR antagonism significantly reduced sociability Z-scores in saline-treated males (*p* = 0.0168), an effect absent in psilocybin-treated males receiving the same antagonist (**Figure 3G**), which was mirrored in reduced sociability indices (*p* = 0.0077; **Figure 3H**). Although no significant 5-HTR antagonism effects were evident in psilocybin-treated mice, 5-HT1AR antagonist pre-treatment showed a trend toward reduced sociability (*p* = 0.0584; **Figure 3H**). At 24 h, 5-HTR antagonism did not affect overall locomotor activity or social preference in males (**Supplementary Data 6**) or females (**Supplementary Data 7**). At 7 d post-administration, psilocybin significantly reduced the sociability Z-score (*p* = 0.0023) in females, reflecting a shift in familiarity preference without affecting other measures. In males, psilocybin increased grooming 7 d following treatment (*p* = 0.0363; **Supplementary Data 8**), opposing 4 h results.

**Figure 3.**
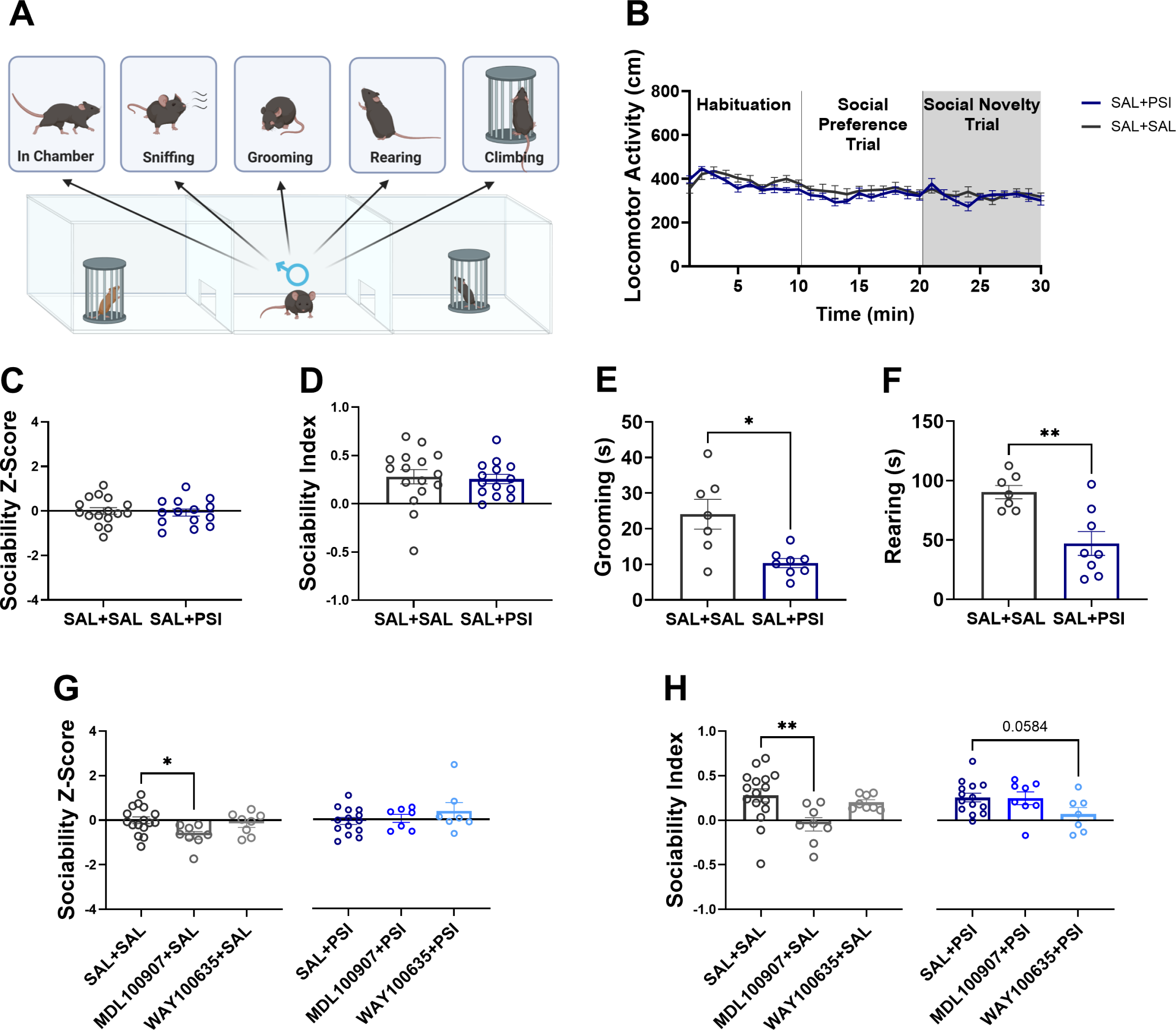
Psilocybin does not affect sociability but reduces grooming and rearing behaviour in male mice 24 hours after administration. **(A)** Schematic of the social novelty trial and the behaviours scored. **(B)** No effects of psilocybin were seen in locomotor activity of mice across the duration of the experiment. **(C,D)** Psilocybin did not affect the sociability of mice. **(E)** Psilocybin significantly increased grooming behaviour (*p* = 0.0159) and it **(F)** significantly reduced rearing behaviour (*p* = 0.0033). **(G)** In SAL-treated mice, 5-HT2AR antagonism reduced sociability Z-score (*p* = 0.0168). No effects of 5-HTR antagonism were observed in PSI-treated mice. **(H)** Similarly, 5-HT2AR antagonism reduced sociability in SAL-treated mice (*p* = 0.0077), while 5-HT1AR antagonism in PSI-treated mice showed a trend toward reduced sociability (*p* = 0.0584). Psilocybin (PSI), saline (SAL), 5-HT2AR antagonist (MDL100907), 5-HT1AR antagonist (WAY100635). Data are presented as mean ± SEM. Statistical analyses were performed using unpaired t-test, one-way ANOVA or two-way ANOVA with Šidák post hoc tests. *p< 0.05, **p< 0.01.

### Psilocybin differentially modulates social motivation in male and female mice at 24 hours

Since 3-chamber sociability is driven by motivational processes, we assessed the effects of psilocybin on social motivation using a barrier climbing test, observing distinct responses in males and females 24 h following administration of saline or psilocybin (with or without antagonism of the 5-HTR subtypes; **Figure 4A**). In females, psilocybin did not alter locomotor activity (**Figure 4B**) or the latency to cross the barrier across different trial phases (**Figure 4C**). However, psilocybin-treated females showed a trend toward increased social motivation, as indicated by a higher sociability Z-score for a novel conspecific (*p* = 0.0566; **Figure 4D**). Latency to cross was unaffected by 5-HTR antagonism in either treatment group (**Figure 4E**), whereas psilocybin-elevated sociability was blocked by 5-HT2AR but not 5-HT1AR antagonism (*p* = 0.0694; **Figure 4F**).

**Figure 4.**
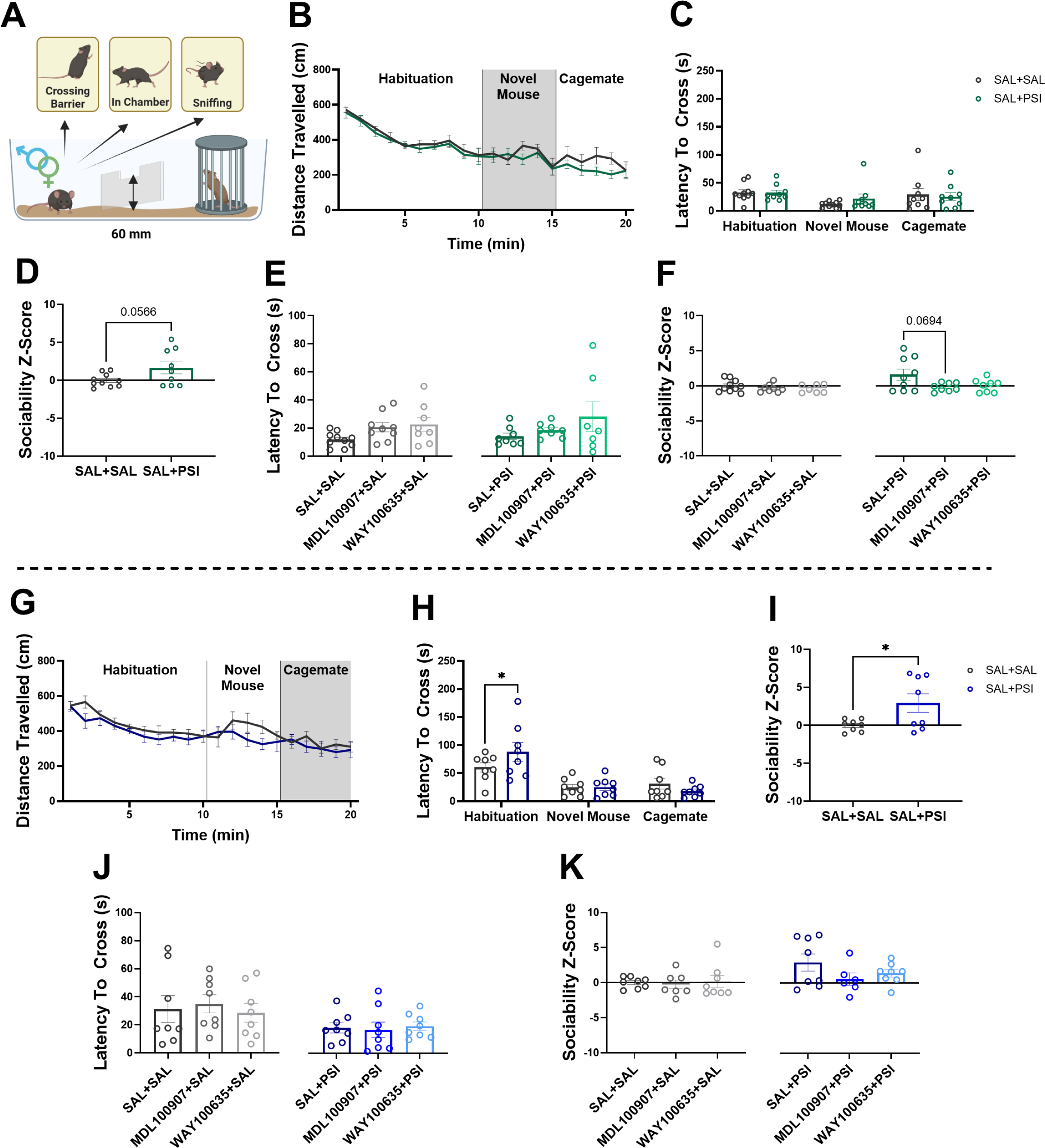
Sex-specific motivational shifts in social preference emerge 24 hours after psilocybin administration. **(A)** Schematic representation of the barrier climbing test and the behavioural measures assessed. **(B)** Locomotor activity and **(C)** the latency to cross the barrier across the trial was not affected by psilocybin in female mice. **(D)** Psilocybin increased the trend towards increased preference towards novelty in female mice (*p* = 0.0566). **(E)** 5-HTR antagonism did not significantly alter barrier crossing latency in female mice and **(F)** their sociability Z-score when exploring the novel conspecific however, in PSI-treated mice, 5-HT2AR antagonism showed a trend toward reducing sociability (*p* = 0.0694). **(G)** In male mice, no effects of psilocybin were seen in locomotor activity across the duration of the experiment. **(H)** Psilocybin increased latency to cross the barrier during the habituation phase (*p* = 0.0407). **(I)** PSI-treated male mice exhibited significantly higher sociability Z-score toward their cage mates compared to the controls (*p* = 0.0349). **(J)** 5-HTR antagonism did not significantly affect latency to cross or **(K)** sociability of male mice when exploring their cage mates. Female mice (A-E), male mice (F-J), psilocybin (PSI), saline (SAL), 5-HT2AR antagonist (MDL100907), 5-HT1AR antagonist (WAY100635). Data are presented as mean ± SEM. Statistical analyses were performed using unpaired t-test, one-way ANOVA or two-way ANOVA with Šidák post hoc tests. *p< 0.05.

In males, psilocybin had no effect on overall locomotor activity (**Figure 4G**) but significantly increased the latency to cross the barrier upon arena introduction (*p* = 0.0407; **Figure 4H**) and enhanced social preference toward (familiar) cage mates (*p* = 0.0349; **Figure 4I**). Neither social motivation nor sociability in male mice was influenced by 5-HTR antagonism under control or psilocybin-treated conditions (**Figure 4J, K**) and 5-HT receptor antagonism did not affect locomotion or sociability Z-scores in either sex on this task (**Supplementary Data 9**).

### Psilocybin prolongs nucleus accumbens (NAc) dopamine release in females during initial familiar social interaction at 7 days post administration and reduces immediate dopaminergic response in males

Based on the dopamine (DA) system’s critical role in social motivation and exploratory behaviour, we examined DA dynamics in the NAc during barrier climbing using fiber photometry to record GRAB_DA_-mediated fluorescence as a proxy for DA release (**Figure 5A**). Since repeated interactions can attenuate DA release sensitivity, we focused our analysis on DA release during the first social interaction. Striatal DA was monitored in male and female mice 24 h after administration of either psilocybin or saline, during social exploration involving an empty cage, a familiar or novel mouse (**Figure 5C** for females; **Figure 5G** for males). DA dynamics showed sex-and time-dependent effects. In females, psilocybin did not significantly affect mean or peak DA release during the first 2 s after sniffing familiar or novel mice (**Figure 5D**). In males, psilocybin significantly reduced peak DA release during initial contact (sniffing) with a novel mouse (*p* = 0.0386), although overall and immediate average DA release were not significantly altered (**Figure 5H**). At 7 d, females exhibited prolonged DA release elevation during the first interaction with the familiar mouse, but not the novel one (**Figure 5E**). though not reflected in the immediate (0-2 s) mean or peak DA release (**Figure 5F**). In males, psilocybin did not affect total DA release at 7 d (**Figure 5I**) but again significantly reduced immediate peak DA release during the first novel mouse sniff (*p* = 0.0326). Consistent with the view that novel conspecific interactions are more rewarding that familiar ones, saline-treated males showed significantly greater immediate peak DA release when sniffing a novel mouse compared to a familiar one (*p* = 0.0353; **Figure 5J**), which was absent following psilocybin treatment.

**Figure 5.**
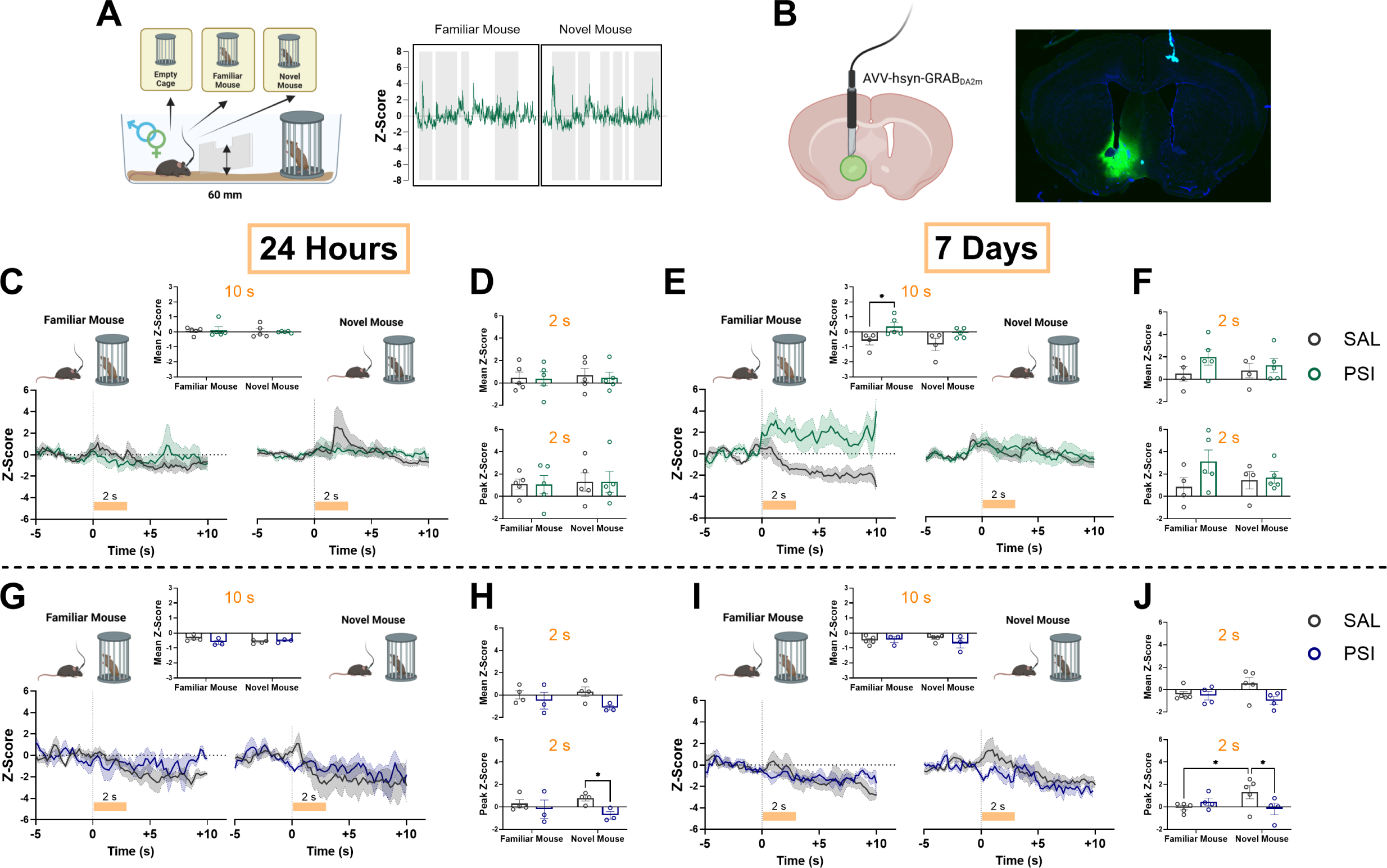
Psilocybin enhances NAc dopamine (DA) release in female mice when sniffing a familiar conspecific at 7 days post-administration and suppresses novelty-evoked dopamine in males at 24 hours and 7 days during the barrier climbing test. **(A)** Schematic of the experimental design and the average GRAB_DA2m_ signal across the full trial when the mouse was in the social zone (shaded area) versus the non-social zone. **(B)** Representative Fiber placement and virus injection in the NAc. **(C)** Representative GRAB_DA2m_ traces from the first interaction with a familiar or a novel mouse at 24 hours post-administration of PSI or SAL control in female mice. Inset: no significant effect of PSI on DA release were observed during the first interaction with either mouse. **(D)** At 24 hours in females, PSI did not alter DA release within the first 2 seconds of sniffing either mouse. **(E)** At 7 days post-administration, representative traces show prolonged DA release in response to familiar (but not novel) sniffing in PSI-treated female mice. Inset: PSI significantly increased DA release within the first 10 seconds of familiar sniffing (*p* = 0.0317), **(F)** but early DA levels (0-2 s) remained unchanged. **(G)** GRAB_DA2m_ traces recorded in male mice during the first sniff of a familiar or novel conspecific, 24 hours after PSI or SAL treatment. Inset: No significant effects of PSI on DA release were observed within the first 10 seconds of the interaction. **(H)** At 24 hours, PSI significantly reduced immediate (0–2 s) DA release during novel sniffing in males (*p* = 0.0386), with no effect on familiar sniffing. **(I)** Representative GRAB_DA2m_ signal recordings from male mice during their first interaction of either a familiar or a novel conspecific, 7 days post-administration of PSI or SAL. Inset: No significant differences in total DA release within 10 seconds post-interaction. **(J)** At 7 days, PSI significantly reduced immediate novelty-evoked DA release in males (*p* = 0.0326); SAL-treated males showed greater DA release to novel vs familiar sniffing (*p* = 0.0353). Female mice (C-F), male mice (G-J), psilocybin (PSI), saline (SAL) and data are presented as mean ± SEM. Statistical analyses were performed using unpaired t-test, one-way ANOVA or two-way ANOVA with Fisher’s LSD post hoc tests. *p< 0.05.

## Discussion

This study demonstrates that psilocybin produces sex- and time-specific effects on social behaviour, mediated by distinct serotonergic and dopaminergic mechanisms. Rather than inducing uniform “prosocial” outcomes, psilocybin modulated behaviour in a context- and sex-dependent manner, even in unstressed, socially housed mice, addressing a key gap in preclinical psychedelic research, where sex differences remain underexplored. We examined acute and delayed effects of a single psilocybin dose on multiple aspects of sociability in male and female mice and assessed the involvement of 5-HT1AR and 5-HT2AR. We also measured DA release in the NAc, a key region within social reward circuits. Given the established interplay between serotonin and DA in regulating cognition and emotion [59], our findings contribute important insights into the neurobiological substrates underlying psychedelic action.

### Acute prosocial behaviour and temperature regulation

Within 60 min of psilocybin administration, female mice exhibited pronounced increases in huddling behaviour, initially hypothesised to reflect thermoregulatory social bonding [60]. The reduction in core body temperature observed in our study has previously been reported only at higher doses of psilocybin (3 and 10 mg/kg), not at lower doses (0.03–1 mg/kg), with no sex differences reported [61]. This discrepancy likely arises from methodological differences: we continuously monitored temperature at 5-min intervals for 1 h post-administration using an automated system, allowing detection of transient fluctuations, whereas the previous study assessed temperature only once at 30 min, potentially overlooking dynamic thermoregulatory responses. Crucially, the psilocybin-induced increase in huddling was abolished by both 5-HT1AR and 5-HT2AR antagonists, but this was not recapitulated in temperature measurements, indicating a more complex interplay between serotonergic signalling and temperature regulation. Specifically, 5-HT1AR antagonism reduced body temperature and suppressed huddling, pointing to its role in tonic thermoregulation and social bonding. In contrast, 5-HT2AR antagonism only reduced temperature when combined with psilocybin and partially disrupted huddling, suggesting it modulates the temporal dynamics of social behaviour rather than thermoregulation *per se*. At 4 h, 5-HT2AR antagonism blocked psilocybin-induced sociability without affecting locomotion, whereas 5-HT1AR effects were confounded by reduced activity. Together, these findings suggest a dissociation between hypothermia and social bonding, with 5-HT1AR playing a more central role, consistent with previous findings [62].

Male mice did not exhibit psilocybin-induced huddling or changes in body temperature, consistent with reports that females are more sensitive to the acute behavioural effects of psilocybin [63]. Interestingly, both 5-HT1AR and 5-HT2AR antagonists reduced aggression in males, an effect reversed by psilocybin co-administration. This suggests psilocybin engages broader serotonergic circuits that override receptor-specific modulation of aggression, contrasting with previous studies where 5-HT1AR agonists suppressed aggression and antagonism reversed these effects [62]. These differing outcomes may reflect compensatory signalling or altered serotonergic tone in pair-housed animals compared to the socially isolated animals in earlier work.

Sex differences in serotonergic architecture may further explain these divergent effects. Women exhibit higher 5-HT1AR binding potential in multiple brain regions, though binding fluctuates with the menstrual cycle [64, 65], and this receptor is a central target of SSRIs in treating social anxiety and depression [66–68]. In contrast, men show higher cortical 5-HT2AR expression [64]. Similar patterns are observed in rodents, with female rats exhibiting higher 5-HT1AR signalling and males showing greater cortical 5-HT2AR expression [69], potentially explaining the differential involvement of these receptors in social and stereotypic behaviours in our study.

Psilocybin-induced hypothermia persisted even when 5-HT1AR or 5-HT2AR were antagonised, suggesting alternative mechanisms. A strong candidate is the 5-HT7R, which is implicated in thermoregulation and is widely expressed in the preoptic area (POA) of the hypothalamus, raphe nuclei, and parabrachial nucleus, all of which are central to autonomic temperature control [70, 71]. Notably, 5-HT7R agonism induces hypothermia, and increased serotonergic tone following psilocybin administration may recruit this receptor [72–74]. Additionally, GABA-ergic and glutamatergic signalling within the POA may interact with serotonergic inputs to shape psilocybin’s hypothermic effects [75, 76].

### Social preference and exploration

Consistent with previous reports [36, 77], psilocybin did not affect social preference in either sex at 24 h or 7 d post-administration. However, at 4 h, female mice showed increased grooming and social exploration, behaviours associated with both heightened anxiety and sociability in C57BL/6 mice [78]. These prosocial effects were abolished by both 5-HT1AR and 5-HT2AR antagonism, confirming involvement of these receptor subtypes. At 24 h, psilocybin reduced grooming and rearing in male mice, behaviours that may reflect anxiolytic effects or reduced behavioural responsiveness. Rearing, more frequent in males [79], is reduced by psilocin through 5-HT2AR mechanisms [80], while grooming is suppressed by SSRIs, 5-HT1AR agonists [81] and high doses of psilocybin or LSD [82–84].

Although earlier findings reported increased sociability in stressed female mice 24 h after psilocybin treatment [85], we did not observe similar prosocial effects at this time point in our unstressed mice. However, robust effects at other time points suggest that stress or social deficits are not necessary preconditions for psilocybin’s behavioural efficacy. While prior work showed that psilocybin can reverse social impairments in autism models without affecting controls [77], our data indicate that psilocybin can modulate social behaviour in healthy animals in a time- and context-dependent manner.

Our data support previous evidence that 5-HT1AR plays a key role in mediating social effects of serotonergic compounds. WAY-100635 blocked psilocybin-induced prosocial behaviours, consistent with reports showing this antagonist prevented MDMA-induced increases in social interaction in male rats [86]. These findings suggest psilocybin initiates dynamic processes that unfold over time and differ between sexes. The persistence of effects beyond the acute pharmacological window may reflect underlying mechanisms such as neuroplastic changes, altered social learning, or sustained serotonergic adaptations. The female-specific shift toward familiar social targets at later time points may indicate memory consolidation processes. Time-dependent effects could also be influenced by interactions between serotonergic and estrogen signalling [48]. Although we did not control for the estrous cycle in all behavioural tests, no significant correlation was observed between estrous stage and sociability in female mice (see **Supplementary Methods** & **Supplementary Data 10**). However, studies in female rats show that those in the estrous and proestrous phases (when estradiol and progesterone are highest) were less affected by psilocin and LSD compared to male rats [87, 88].

### Dopamine dynamics and social reward salience

NAc DA dynamics revealed a possible mechanistic substrate for sex-specific behavioural responses. In females, psilocybin did not alter acute DA responses but induced a prolonged elevation in DA release during familiar, but not novel, interactions at 7 d post-administration. This may reflect enhanced social memory or encoding of familiar cues, consistent with findings that familiarity plays a larger role in female prosocial [89]. The psilocybin-induced increase in NAc DA during familiar interactions may reflect a recalibration of social reward circuits, wherein familiar social experiences become re-evaluated as uniquely rewarding within a new affective or cognitive context. The delayed response aligns with neuroplastic adaptations, such as increased spine density in the female mouse cortex post-psilocybin [90] and improved cognitive flexibility [91, 92]. These interactions underscore that NAc DA signals multiple dimensions including salience, motivation, valence, prediction error. Psilocybin may recalibrate these behavioural computations in sex-specific ways.

In males, psilocybin reduced peak DA release during novel social encounters at both 24 h and 7 d, aligning with behavioural shifts toward familiar preference and reduced novelty salience. Saline-treated controls displayed the expected novelty-biased DA response [93], which was absent in psilocybin-treated males, suggesting psilocybin dampens novelty-driven dopaminergic activity, potentially reshaping the social reward landscape. Whether this reflects reduced novelty “appeal” or increased reward value of familiar conspecifics remains uncertain. However, since the photometry-paired task did not require choosing between familiar and novel conspecifics, the latter explanation may be more plausible. Consistent with our observations, a recent study reported sex-specific responses, with psilocybin eliciting stronger behavioural effects and calcium transients in the paraventricular nucleus of female mice compared to males [94].

### Limitations and future directions

Some limitations warrant consideration: First, we only tested a single psilocybin dose, limiting conclusions about dose-response relationships. Second, our study focused on 5-HT1AR and 5-HT2AR, while other receptors like the 5-HT7R, 5-HT1BR and 5-HT2CR that regulate thermoregulation and social behaviour should be explored [71, 95]. We observed that 5-HT1AR antagonism reduced locomotor activity in female mice at 4 h, potentially reflecting sex-specific sensitivity to the antagonist dose, which complicates interpretation of the associated sociability findings. While this confound should be acknowledged, effects across other measures still offer preliminary insights into receptor-specific contributions warranting further investigation. Additionally, although we discuss serotonergic-dopaminergic interactions, we did not assess the oxytocin system, which plays a central role in social recognition and bonding [96]. Finally, while fiber photometry is a powerful tool for monitoring DA dynamics, it captures only population-level changes and does not quantify absolute concentrations or resolve fine-scale spatial activity. These recordings do not clarify whether psilocybin’s effects arise from direct modulation of DA neuron activity in the midbrain or from local changes in the NAc, potentially mediated by 5-HT2AR signalling [97, 98].

Despite these limitations, this study is the first, to our knowledge, to assess acute and delayed effects of psilocybin on multiple dimensions of social behaviour and DA release in male and female mice. The single-dose paradigm offers clearer interpretation, avoiding confounds from repeated administration and reflecting current clinical contexts. The focus on 5-HT1AR and 5-HT2AR, two of the most well-characterised 5-HT receptors, strengthens interpretability. We demonstrate that psilocybin modulates social behaviour and core body temperature through sexually dimorphic and temporally dynamic mechanisms, mediated by 5-HT1AR and 5-HT2AR signalling and NAc dopaminergic pathways. While 5-HT2AR has historically dominated psychedelic research, our results highlight the need to broaden this focus. These findings challenge assumptions of universal prosocial effects and underscore the critical importance of incorporating SABV in preclinical models and therapeutic development. Future studies should investigate broader receptor systems, hormonal influences, and circuit-level mechanisms to further elucidate the complex, sex-specific neurobiology underlying psychedelic modulation of social cognition and behaviour.

## Author Contributions

SS led the experimental work and wrote the initial manuscript draft. SS, KM, LO & EG contributed to the acquisition and analysis of data for the work. KC, ZBA & CJF contributed to the analysis and interpretation of data and critical review of the manuscript drafts. CJF conceptualised the study and supervised the authorship team. All authors approved of the submitted version of the manuscript, and agree to be accountable for all aspects of data contained within.

## Acknowledgements

We acknowledge the use of Biorender.com for elements of the figures and the AI tool claude.ai for assistance scanning the final manuscript for redundant text to facilitate adherence to the word limit.

## Funding

This work was supported by a National Health and Medical Research Council (NHMRC) of Australia Ideas Grant (GNT2011334) awarded to CJF. SS is supported by a Monash Biomedicine Postgraduate Discovery Scholarship (Mbio).

## Competing Interests

The authors have nothing to disclose.

## Notes

### Competing Interest Statement

The authors have declared no competing interest.

### Summary of Updates

Streamlined text to reduce redundancy and adhere to journal submission word limits.

## References

1. Dunbar, R.I.M., The Anatomy of Friendship. Trends in Cognitive Sciences, 2018. 22(1): p. 32–51.

2. Brotman, M.A., et al., Irritability in Youths: A Translational Model. Am J Psychiatry, 2017. 174(6): p. 520–532.

3. Kennedy, D.P. and R. Adolphs, The social brain in psychiatric and neurological disorders. Trends Cogn Sci, 2012. 16(11): p. 559–72.

4. Preller, K.H. and F.X. Vollenweider, Modulation of Social Cognition via Hallucinogens and “Entactogens”. Front Psychiatry, 2019. 10: p. 881.

5. Rodríguez Arce, J.M. and M.J. Winkelman, Psychedelics, Sociality, and Human Evolution. Front Psychol, 2021. 12: p. 729425.

6. Peck, S.K., et al., Psilocybin therapy for females with anorexia nervosa: a phase 1, open-label feasibility study. Nat Med, 2023. 29(8): p. 1947–1953.

7. Barber, G.S. and S.T. Aaronson, The Emerging Field of Psychedelic Psychotherapy. Curr Psychiatry Rep, 2022. 24(10): p. 583–590.

8. Carhart-Harris, R.L. and G.M. Goodwin, The Therapeutic Potential of Psychedelic Drugs: Past, Present, and Future. Neuropsychopharmacology, 2017. 42(11): p. 2105–2113.

9. Carhart-Harris, R.L., et al., Psilocybin for treatment-resistant depression: fMRI-measured brain mechanisms. Scientific reports, 2017. 7(1): p. 1–11.

10. Davis, A.K., F.S. Barrett, and R.R. Griffiths, Psychological flexibility mediates the relations between acute psychedelic effects and subjective decreases in depression and anxiety. Journal of Contextual Behavioral Science, 2020. 15: p. 39–45.

11. Davis, A.K., et al., Effects of Psilocybin-Assisted Therapy on Major Depressive Disorder: A Randomized Clinical Trial. JAMA Psychiatry, 2021. 78(5): p. 481–489.

12. Griffiths, R.R., et al., Survey of subjective “God encounter experiences”: Comparisons among naturally occurring experiences and those occasioned by the classic psychedelics psilocybin, LSD, ayahuasca, or DMT. PLoS One, 2019. 14(4): p. e0214377.

13. Moreno, F.A., et al., Safety, tolerability, and efficacy of psilocybin in 9 patients with obsessive-compulsive disorder. J Clin Psychiatry, 2006. 67(11): p. 1735–40.

14. Ross, S., et al., Rapid and sustained symptom reduction following psilocybin treatment for anxiety and depression in patients with life-threatening cancer: a randomized controlled trial. Journal of psychopharmacology, 2016. 30(12): p. 1165–1180.

15. Forstmann, M., et al., Transformative experience and social connectedness mediate the mood-enhancing effects of psychedelic use in naturalistic settings. Proc Natl Acad Sci U S A, 2020. 117(5): p. 2338–2346.

16. Pokorny, T., et al., Effect of Psilocybin on Empathy and Moral Decision-Making. Int J Neuropsychopharmacol, 2017. 20(9): p. 747–757.

17. Preller, K.H., et al., Effects of serotonin 2A/1A receptor stimulation on social exclusion processing. Proc Natl Acad Sci U S A, 2016. 113(18): p. 5119–24.

18. Dolder, P.C., et al., LSD Acutely Impairs Fear Recognition and Enhances Emotional Empathy and Sociality. Neuropsychopharmacology, 2016. 41(11): p. 2638–46.

19. Rucker, J.J., et al., The effects of psilocybin on cognitive and emotional functions in healthy participants: Results from a phase 1, randomised, placebo-controlled trial involving simultaneous psilocybin administration and preparation. J Psychopharmacol, 2022. 36(1): p. 114–125.

20. Bhatt, K.V. and C.R. Weissman, The effect of psilocybin on empathy and prosocial behavior: a proposed mechanism for enduring antidepressant effects. Npj Ment Health Res, 2024. 3(1): p. 7.

21. Jungwirth, J., et al., Psilocybin increases emotional empathy in patients with major depression. Molecular Psychiatry, 2024.

22. Carhart-Harris, R.L., et al., Psychedelics and connectedness. Psychopharmacology (Berl), 2018. 235(2): p. 547–550.

23. Griffiths, R., et al., Mystical-type experiences occasioned by psilocybin mediate the attribution of personal meaning and spiritual significance 14 months later. J Psychopharmacol, 2008. 22(6): p. 621–32.

24. Griffiths, R.R., et al., Psilocybin occasioned mystical-type experiences: immediate and persisting dose-related effects. Psychopharmacology (Berl), 2011. 218(4): p. 649–65.

25. McCulloch, D.E., et al., Psilocybin-Induced Mystical-Type Experiences are Related to Persisting Positive Effects: A Quantitative and Qualitative Report. Front Pharmacol, 2022. 13: p. 841648.

26. Aleksandrova, L.R. and A.G. Phillips, Neuroplasticity as a convergent mechanism of ketamine and classical psychedelics. Trends in Pharmacological Sciences, 2021. 42(11): p. 929–942.

27. van Elk, M. and D.B. Yaden, Pharmacological, neural, and psychological mechanisms underlying psychedelics: A critical review. Neuroscience & Biobehavioral Reviews, 2022. 140: p. 104793.

28. Gabay, A.S., et al., Psilocybin and MDMA reduce costly punishment in the Ultimatum Game. Sci Rep, 2018. 8(1): p. 8236.

29. Mason, N.L., et al., Sub-Acute Effects of Psilocybin on Empathy, Creative Thinking, and Subjective Well-Being. J Psychoactive Drugs, 2019. 51(2): p. 123–134.

30. Kolasa, M., et al., Unraveling psilocybin’s therapeutic potential: behavioral and neuroplasticity insights in Wistar-Kyoto and Wistar male rat models of treatment-resistant depression. Psychopharmacology (Berl), 2025. 242(7): p. 1607–1625.

31. De Gregorio, D., et al., Lysergic acid diethylamide (LSD) promotes social behavior through mTORC1 in the excitatory neurotransmission. Proc Natl Acad Sci U S A, 2021. 118(5).

32. Gattuso, J.J., et al., Chronic psilocybin administration increases sociability and alters the gut microbiome in male wild-type mice but not in a preclinical model of obsessive-compulsive disorder. Neuropharmacology, 2025. 279: p. 110648.

33. Markopoulos, A., et al., Lysergic acid diethylamide (LSD) promotes social behaviour through 5-HT2A and ampa in the medial prefrontal cortex (MPFC). European Psychiatry, 2021. 64(S1): p. S416–S417.

34. Song, C., et al., Lasting effect of psilocybin on sociability can be blocked by DNA methyltransferase inhibition. bioRxiv, 2025.

35. Nardou, R., et al., Psychedelics reopen the social reward learning critical period. Nature, 2023. 618(7966): p. 790–798.

36. Lu, O.D., et al., A multi-institutional investigation of psilocybin’s effects on mouse behavior. bioRxiv, 2025: p. 2025.04.08.647810.

37. Duerler, P., F.X. Vollenweider, and K.H. Preller, A neurobiological perspective on social influence: Serotonin and social adaptation. Journal of Neurochemistry, 2022. 162(1): p. 60–79.

38. Ko, J., Neuroanatomical Substrates of Rodent Social Behavior: The Medial Prefrontal Cortex and Its Projection Patterns. Frontiers in Neural Circuits, 2017. 11.

39. Luo, J., The Neural Basis of and a Common Neural Circuitry in Different Types of Pro-social Behavior. Frontiers in Psychology, 2018. 9.

40. Kim, B.S., et al., Differential regulation of observational fear and neural oscillations by serotonin and dopamine in the mouse anterior cingulate cortex. Psychopharmacology, 2014. 231: p. 4371–4381.

41. Yamagishi, A., J. Lee, and N. Sato, Oxytocin in the anterior cingulate cortex is involved in helping behaviour. Behavioural Brain Research, 2020. 393: p. 112790.

42. Lockwood, P.L. and M.K. Wittmann, Ventral anterior cingulate cortex and social decision-making. Neuroscience & Biobehavioral Reviews, 2018. 92: p. 187–191.

43. Sladky, R., et al., Disrupted Effective Connectivity Between the Amygdala and Orbitofrontal Cortex in Social Anxiety Disorder During Emotion Discrimination Revealed by Dynamic Causal Modeling for fMRI. Cerebral Cortex, 2013. 25(4): p. 895–903.

44. Liu, Y. and Z. Wang, Nucleus accumbens oxytocin and dopamine interact to regulate pair bond formation in female prairie voles. Neuroscience, 2003. 121(3): p. 537–544.

45. Nardou, R., et al., Oxytocin-dependent reopening of a social reward learning critical period with MDMA. Nature, 2019. 569(7754): p. 116–120.

46. Effinger, D.P., et al., Increased reactivity of the paraventricular nucleus of the hypothalamus and decreased threat responding in male rats following psilocin administration. Nat Commun, 2024. 15(1): p. 5321.

47. Bangasser, D.A. and A. Cuarenta, Sex differences in anxiety and depression: circuits and mechanisms. Nat Rev Neurosci, 2021. 22(11): p. 674–684.

48. Shadani, S., et al., Potential Differences in Psychedelic Actions Based on Biological Sex. Endocrinology, 2024. 165(8).

49. Kuehner, C., Why is depression more common among women than among men? Lancet Psychiatry, 2017. 4(2): p. 146–158.

50. Christiansen, D.M. and E.T. Berke, Gender- and Sex-Based Contributors to Sex Differences in PTSD. Curr Psychiatry Rep, 2020. 22(4): p. 19.

51. Farhane-Medina, N.Z., et al., Factors associated with gender and sex differences in anxiety prevalence and comorbidity: A systematic review. Sci Prog, 2022. 105(4): p. 368504221135469.

52. Timko, C.A., L. DeFilipp, and A. Dakanalis, Sex Differences in Adolescent Anorexia and Bulimia Nervosa: Beyond the Signs and Symptoms. Curr Psychiatry Rep, 2019. 21(1): p. 1.

53. Clayton, J.A., Applying the new SABV (sex as a biological variable) policy to research and clinical care. Physiology & Behavior, 2018. 187: p. 2–5.

54. Conn, K., et al., Psilocybin restrains activity-based anorexia in female rats by enhancing cognitive flexibility: contributions from 5-HT1A and 5-HT2A receptor mechanisms. Mol Psychiatry, 2024. 29(10): p. 3291–3304.

55. Guilloux, J.P., et al., Integrated behavioral z-scoring increases the sensitivity and reliability of behavioral phenotyping in mice: relevance to emotionality and sex. J Neurosci Methods, 2011. 197(1): p. 21–31.

56. Kohl, J., et al., Functional circuit architecture underlying parental behaviour. Nature, 2018. 556(7701): p. 326–331.

57. Salais-López, H., et al., Maternal Motivation: Exploring the Roles of Prolactin and Pup Stimuli. Neuroendocrinology, 2021. 111(9): p. 805–830.

58. Swart, J.M., et al., Changes in maternal motivation across reproductive states in mice: A role for prolactin receptor activation on GABA neurons. Horm Behav, 2021. 135: p. 105041.

59. McCoy, K., et al., Separate or inseparable? Serotonin and dopamine system interactions may underlie the therapeutic potential of psilocybin for anorexia nervosa. Physiology & Behavior, 2025. 298: p. 114957.

60. Landen, J.G., et al., Huddling substates in mice facilitate dynamic changes in body temperature and are modulated by Shank3b and Trpm8 mutation. Commun Biol, 2024. 7(1): p. 1186.

61. McGriff, S.A., et al., Psychedelic-like effects induced by 2,5-dimethoxy-4-iodoamphetamine, lysergic acid diethylamide, and psilocybin in male and female C57BL/6J mice. Psychopharmacology, 2025. 242(10): p. 2249–2260.

62. Tan, O., L.J. Martin, and M.T. Bowen, Divergent pathways mediate 5-HT1A receptor agonist effects on close social interaction, grooming and aggressive behaviour in mice: Exploring the involvement of the oxytocin and vasopressin systems. Journal of Psychopharmacology, 2020. 34(7): p. 795–805.

63. Farinha-Ferreira, M., et al., Concurrent stress modulates the acute and post-acute effects of psilocybin in a sex-dependent manner. Neuropharmacology, 2025. 266: p. 110280.

64. Moses-Kolko, E.L., et al., *Age, Sex,* and Reproductive Hormone Effects on Brain Serotonin-1A and Serotonin-2A Receptor Binding in a Healthy Population. Neuropsychopharmacology, 2011. 36(13): p. 2729–2740.

65. Jovanovic, H., et al., A PET study of 5-HT1A receptors at different phases of the menstrual cycle in women with premenstrual dysphoria. Psychiatry Res, 2006. 148(2-3): p. 185–93.

66. Martin, V., et al., Key role of the 5-HT1A receptor addressing protein Yif1B in serotonin neurotransmission and SSRI treatment. J Psychiatry Neurosci, 2020. 45(5): p. 344–355.

67. Lanzenberger, R.R., et al., Reduced serotonin-1A receptor binding in social anxiety disorder. Biological psychiatry, 2007. 61(9): p. 1081–1089.

68. Le Poul, E., et al., Early desensitization of somato-dendritic 5-HT1A autoreceptors in rats treated with fluoxetine or paroxetine. Naunyn-Schmiedeberg’s archives of pharmacology, 1995. 352(2): p. 141–148.

69. Pitychoutis, P.M., et al., *5-HT1A*, *5-HT2A, and 5-HT2C receptor mRNA modulation by antidepressant treatment in the chronic mild stress model of depression: sex differences exposed*. Neuroscience, 2012. 210: p. 152–167.

70. Hedlund, P.B., et al., 8-OH-DPAT acts on both 5-HT1A and 5-HT7 receptors to induce hypothermia in rodents. Eur J Pharmacol, 2004. 487(1-3): p. 125–32.

71. Negoiţă, A.-I., et al., The Serotonergic System and Its Role in Thermoregulation. Physiologia, 2025. 5(4): p. 37.

72. Halberstadt, A.L. and M.A. Geyer, Multiple receptors contribute to the behavioral effects of indoleamine hallucinogens. Neuropharmacology, 2011. 61(3): p. 364–81.

73. Andressen, K.W., et al., The atypical antipsychotics clozapine and olanzapine promote down-regulation and display functional selectivity at human 5-HT7 receptors. Br J Pharmacol, 2015. 172(15): p. 3846–60.

74. Okada, M., et al., Effects of Subchronic Administrations of Vortioxetine, Lurasidone, and Escitalopram on Thalamocortical Glutamatergic Transmission Associated with Serotonin 5-HT7 Receptor. Int J Mol Sci, 2021. 22(3).

75. Ishiwata, T., et al., Changes of body temperature and thermoregulatory responses of freely moving rats during GABAergic pharmacological stimulation to the preoptic area and anterior hypothalamus in several ambient temperatures. Brain Research, 2005. 1048(1): p. 32–40.

76. Sengupta, T., et al., L-glutamate microinjection in the preoptic area increases brain and body temperature in freely moving rats. Neuroreport, 2014. 25(1): p. 28–33.

77. Song, C., et al., Lasting effect of psilocybin on sociability can be blocked by DNA methyltransferase inhibition. bioRxiv, 2025: p. 2025.03.10.642385.

78. An, X.-L., et al., Strain and Sex Differences in Anxiety-Like and Social Behaviors in C57BL/6J and BALB/cJ Mice. Experimental Animals, 2011. 60(2): p. 111–123.

79. Sturman, O., P.-L. Germain, and J. Bohacek, Exploratory rearing: a context- and stress-sensitive behavior recorded in the open-field test. Stress, 2018. 21(5): p. 443–452.

80. Halberstadt, A.L., S.B. Powell, and M.A. Geyer, Role of the 5-HT2A receptor in the locomotor hyperactivity produced by phenylalkylamine hallucinogens in mice. Neuropharmacology, 2013. 70: p. 218–227.

81. Brookshire, B.R. and S.R. Jones, Direct and indirect 5-HT receptor agonists produce gender-specific effects on locomotor and vertical activities in C57 BL/6J mice. Pharmacology Biochemistry and Behavior, 2009. 94(1): p. 194–203.

82. Kyzar, E.J., A.M. Stewart, and A.V. Kalueff, Effects of LSD on grooming behavior in serotonin transporter heterozygous (Sert⁺/⁻) mice. Behav Brain Res, 2016. 296: p. 47–52.

83. Brownstien, M., et al., Striking long-term beneficial effects of single dose psilocybin and psychedelic mushroom extract in the SAPAP3 rodent model of OCD-like excessive self-grooming. Molecular Psychiatry, 2025. 30(3): p. 1172–1183.

84. Bruno, V., et al., High dose of psilocybin induces acute behavioral changes without inducing conditioned place preference in Sprague-Dawley rats. Journal of Psychopharmacology. 0(0): p. 02698811251368361.

85. Meng, C., et al., Structural basis for psilocybin biosynthesis. Nature Communications, 2025. 16(1): p. 2827.

86. Morley, K.C., J.C. Arnold, and I.S. McGregor, Serotonin (1A) receptor involvement in acute 3,4-methylenedioxymethamphetamine (MDMA) facilitation of social interaction in the rat. Progress in Neuro-Psychopharmacology and Biological Psychiatry, 2005. 29(5): p. 648–657.

87. Tylš, F., et al., Sex differences and serotonergic mechanisms in the behavioural effects of psilocin. Behavioural Pharmacology, 2016. 27(4).

88. Pálenícek, T., et al., Sex differences in the effects of N,N-diethyllysergamide (LSD) on behavioural activity and prepulse inhibition. Prog Neuropsychopharmacol Biol Psychiatry, 2010. 34(4): p. 588–96.

89. Misiołek, K., et al., Prosocial behavior, social reward and affective state discrimination in adult male and female mice. Scientific Reports, 2023. 13(1): p. 5583.

90. Shao, L.X., et al., Psilocybin induces rapid and persistent growth of dendritic spines in frontal cortex in vivo. Neuron, 2021. 109(16): p. 2535–2544.e4.

91. Meshkat, S., et al., Impact of psilocybin on cognitive function: A systematic review. Psychiatry Clin Neurosci, 2024. 78(12): p. 744–764.

92. Fisher, E.L., et al., Psilocybin increases optimistic engagement over time: computational modelling of behaviour in rats. Transl Psychiatry, 2024. 14(1): p. 394.

93. Dai, B., et al., Responses and functions of dopamine in nucleus accumbens core during social behaviors. Cell Rep, 2022. 40(8): p. 111246.

94. Cook, S.G., et al., Psilocybin induces sex- and context-specific recruitment of the stress axis. Current Biology, 2025.

95. Wu, X., et al., 5-HT modulation of a medial septal circuit tunes social memory stability. Nature, 2021. 599(7883): p. 96–101.

96. Raam, T., et al., Hippocampal oxytocin receptors are necessary for discrimination of social stimuli. Nature Communications, 2017. 8(1): p. 2001.

97. Yan, Q.-S., Activation of 5-HT2A/2C receptors within the nucleus accumbens increases local dopaminergic transmission. Brain Research Bulletin, 2000. 51(1): p. 75–81.

98. Alex, K.D. and E.A. Pehek, Pharmacologic mechanisms of serotonergic regulation of dopamine neurotransmission. Pharmacology & Therapeutics, 2007. 113(2): p. 296–320.

