## Supplementary for "Psilocybin modulates social behaviour in male and female mice in a time-dependent manner"

**Supplementary Methods (Detailed Protocols)**

**Animals and Housing - Extended Details**

All animals were obtained from the Monash Animal Research Platform (MARP; Clayton, VIC, Australia). Mice were pair-housed in a climate-controlled room (temperature 22-24°C and humidity 30-50%) on a reverse light cycle (lights on at 2000h, off at 0800h) for 7 days before experiments commenced for acclimation. Mice had ad libitum access to food and water, except during behavioural testing windows. Four sex- and age-matched C57Bl/6 mice were used per cohort as novel mice in the 3-chamber and barrier climbing tests and were housed in a separate room under the same conditions as test mice. All experimental procedures adhered to the Australian Code for the Care and Use of Animals for Scientific Purposes.

**Drug Preparation and Administration - Extended Details**

Psilocybin (USONA Institute Investigational Drug Supply Program; Lot# AMS0167) was dissolved in 0.9% NaCl saline. MDL100907 (volinanserin; Sigma-Aldrich, CAS 139290-65-6) and WAY100635 maleate (Tocris Biosciences, CAS 1092679-51-0) were also dissolved in saline. All drugs were delivered intraperitoneally with a 26-gauge needle at an injection volume of 10 ml/kg.

**Acute Core Body Temperature and Activity Monitoring - Extended Details**

The UID Temperature Monitoring System (Unified Information Devices, Kenosha, WI, USA) was used to measure digital biomarkers continuously 24/7 from conscious, unrestrained mice in their home-cage environment. This approach reduced stress and ensured reliable data collection through a temperature-sensitive microchip in an undisturbed setting. Upon arrival, pair-housed mice were subcutaneously implanted with a 2.1 mm x 13 mm microchip which uses RFID technology for animal identification and body temperature monitoring (model UCT-2112), implanted via a specialized injection device.

**3-Chamber Test - Extended Details**

Mice were placed in the central chamber of an apparatus containing three chambers, each with a wire cage at either end, for a 10-minute habituation period. A 1 min inter-trial interval (ITI) followed, during which mice were confined to the middle chamber to minimize experimenter interference. Subsequently, an unfamiliar, age- and sex-matched mouse (Novel 1) was placed in a wire cage in one side chamber, while a novel object in a wire cage occupied the opposite chamber. The test mouse was then allowed to explore for 10 min (social preference trial). After another 1 min ITI, a second unfamiliar age- and sex-matched mouse (Novel 2) replaced the novel object, with Novel 1 remaining as the familiar mouse. The test mouse was observed for another 10 min (social novelty trial). Locomotor activity, interaction frequency, time spent in each chamber, and social preference were measured using video tracking and EthoVision XT software. The Sociability Index measures relative time spent interacting with either a novel conspecific, an empty cage, or a familiar mouse. Stereotypic behaviours assessed included grooming, rearing, cage climbing, and freezing.

**Barrier Climbing Test - Extended Details**

Behaviour was recorded for a total of 20 min, including habituation to the barrier (10 min), exploration of an unfamiliar age- and sex-matched mouse (5 min), and exploration of a cagemate (5 min). The novel mouse and cagemate were placed in a wire cage on the opposite side of the barrier to the test mouse. A barrier climb was defined as the moment when all four paws of the mouse touched the floor on opposite side of the 60 mm transparent barrier. Each instance of the mouse crossing the barrier and placing all four paws on the opposite side was counted as a barrier climb.

**Estrous Cycle Assessment**

Vaginal smears were collected daily at 1000h over a period of 7 d prior to testing and on test days following procedures. A volume of 10 µL of saline was flushed into the vagina, transferred onto a glass slide, and examined as stained preparations using hematoxylin and eosin (H&E). The stage of the estrous cycle was determined and classified as proestrus, estrus, or metestrus/diestrus, based on observed ratios of cornified epithelial, nucleated epithelial, and polymorphonuclear leukocytes [1].

**Surgical and Viral Injection Procedures - Extended Details**

Mice were anesthetized with isoflurane (2-3%; Pharmachem, QLD, Australia) and administered meloxicam (5 mg/kg, Boehringer Ingelheim, Germany) subcutaneously prior to surgery. Each animal was positioned in a stereotaxic frame (Stoelting, IL, USA) on a heating pad maintained at 37.2°C. Mice were unilaterally injected in the right hemisphere with hsyn-GRABDA2m for dopamine recordings. A volume of 300 nL was injected at a rate of 30 nL/min using a 2 μL Hamilton syringe (Model #7002), and the needle was left in place for 5 min post-infusion to allow for diffusion. Optical fibers (RWD, 0.39 NA; length: 4.7 mm; core: 200 nm) were implanted 0.1 mm above the injection site (final DV: -4.1 mm) and secured using bonding agent (GBond, Japan) and light-cured dental cement (G-aenial Universal Flow, GC Dental).

**Fiber Photometry - Extended Details**

Experiments began at least five weeks post-surgery to allow sufficient time for recovery and viral expression. Recordings were conducted using the RWD R821 fiber photometry system, with 470 nm and 410 nm lasers used for the signal and isosbestic control channels, respectively. Data acquisition was performed using RWD software during the barrier climbing task, and behavioural video timestamps were manually aligned to the photometry recordings.

**Data Analysis - Extended Details**

A significance threshold of p < 0.05 was applied, while p < 0.10 was considered indicative of a trend, though not statistically significant. Depending on the data type, number of groups, and comparisons of interest, appropriate statistical methods were used, including two-tailed unpaired t-tests, one-way and two-way analyses of variance (ANOVA) with Sidak's post hoc multiple comparisons and mixed-effects models.

**References**

1.	Nelson, J.F., et al., *A longitudinal study of estrous cyclicity in aging C57BL/6J mice: I. Cycle frequency, length and vaginal cytology.* Biol Reprod, 1982. **27**(2): p. 327-39.

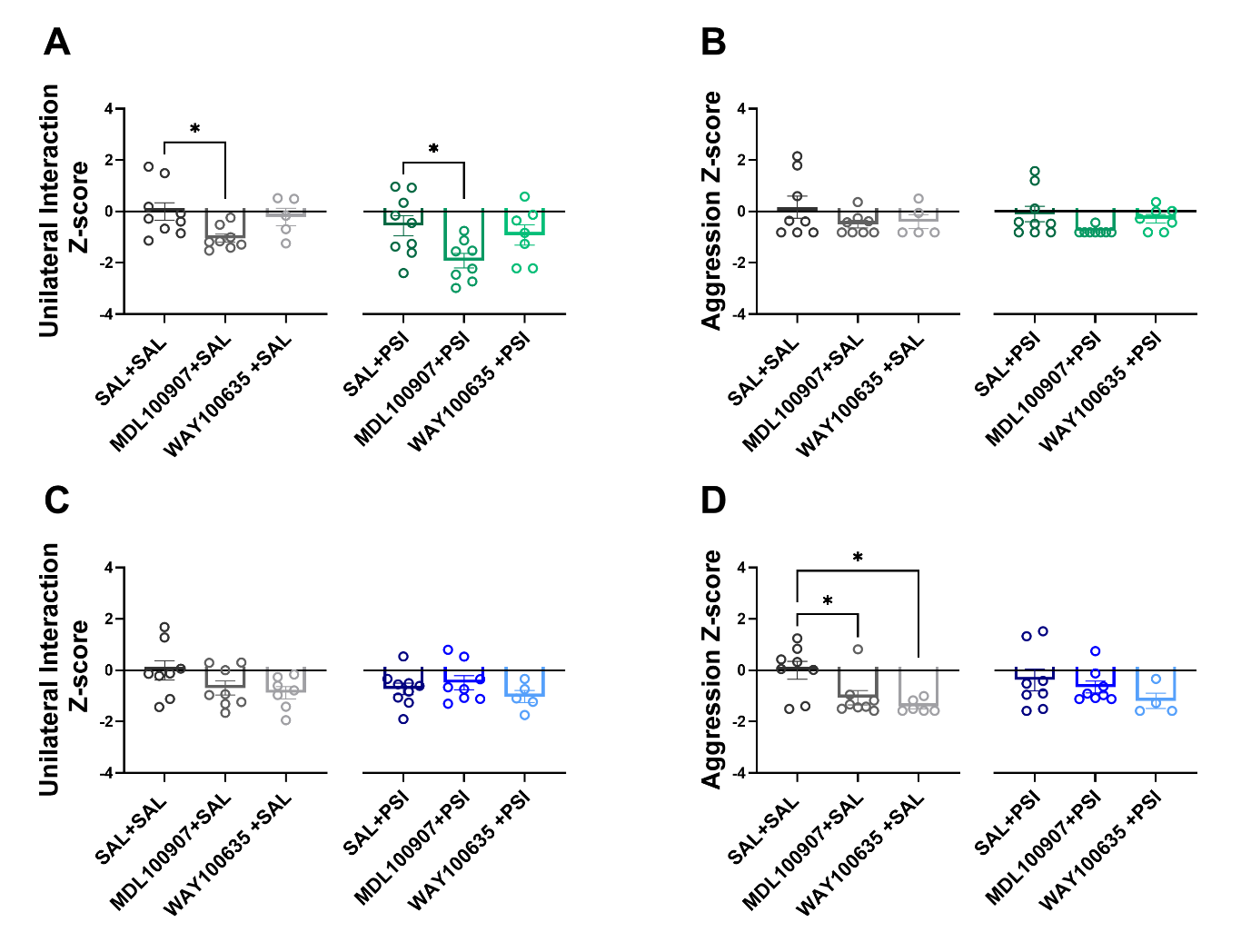

**Supplementary Data 1. 5-HT1AR and 5-HT2AR antagonism reduced unilateral interaction in female mice regardless of psilocybin treatment and decreased aggressive behaviour in male mice only in the absence of psilocybin. (A)** In SAL-treated female mice, 5-HT2AR antagonism significantly reduced the unilateral interaction (*p* = 0.0339) and a similar reduction was observed in PSI-treated females (*p* = 0.0328). **(B)** Neither psilocybin nor 5-HT receptor antagonism significantly affected aggressive behaviour in female mice. **(C)** No significant effects of psilocybin or 5-HT receptor antagonism were observed on unilateral interaction in male mice. **(D)** In SAL-treated male mice, both 5-HT1AR antagonism (*p* = 0.0101) and 5-HT2AR (*p* = 0.0366) antagonism significantly reduced aggressive behaviour. Psilocybin (PSI), saline (SAL), 5-HT2AR antagonist (MDL100907), 5-HT1AR antagonist (WAY100635). Data are presented as mean ± SEM and were analysed using one-way ANOVA with Šidák post hoc tests. *p* < 0.05.

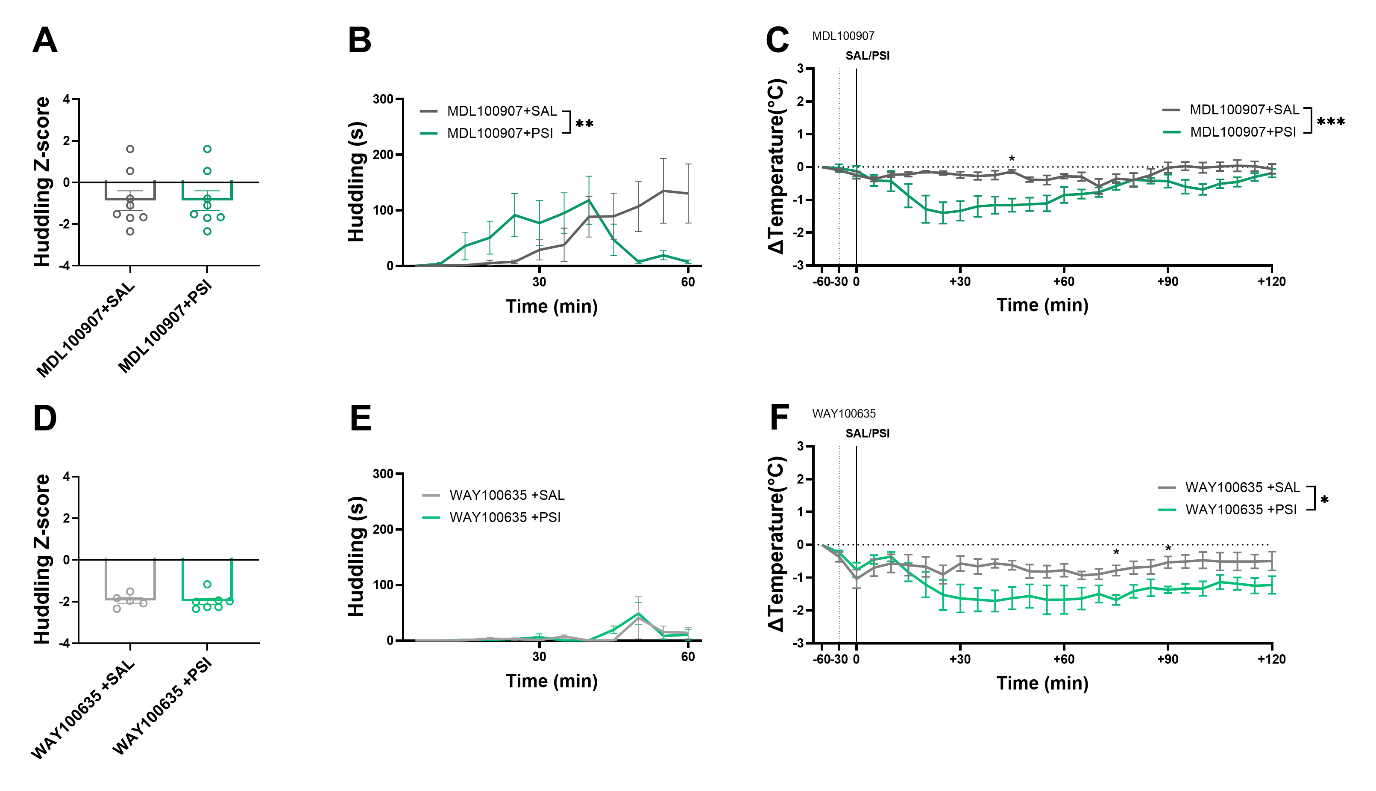

**Supplementary Data 2. 5-HT1AR antagonism reduced acute core body temperature without altering sociability in female mice. (A)** Total huddling behaviour was not affected by PSI in mice pre-treated with the 5-HT2AR antagonist, **(B)** nor was any difference observed throughout the 60-minute trial. **(C)** However, PSI significantly reduced core body temperature in this group, particularly at 45 min post-administration (*p* = 0.0297). **(D)** Similarly, total huddling behaviour was unaffected by PSI in mice pre-treated with the 5-HT1AR antagonist, **(E)** with no changes across the 60-minute observation period. **(F)** In contrast, PSI significantly reduced core body temperature in these mice at 75 min (*p* = 0.0446) and 90 min (*p* = 0.0478) post-administration. Psilocybin (PSI), saline (SAL), 5-HT2AR antagonist (MDL100907), 5-HT1AR antagonist (WAY100635). Data are presented as mean ± SEM. Statistical analyses were performed using unpaired t-test, mixed-effects models or two-way ANOVA with Šidák post hoc tests. **p*< 0.05.

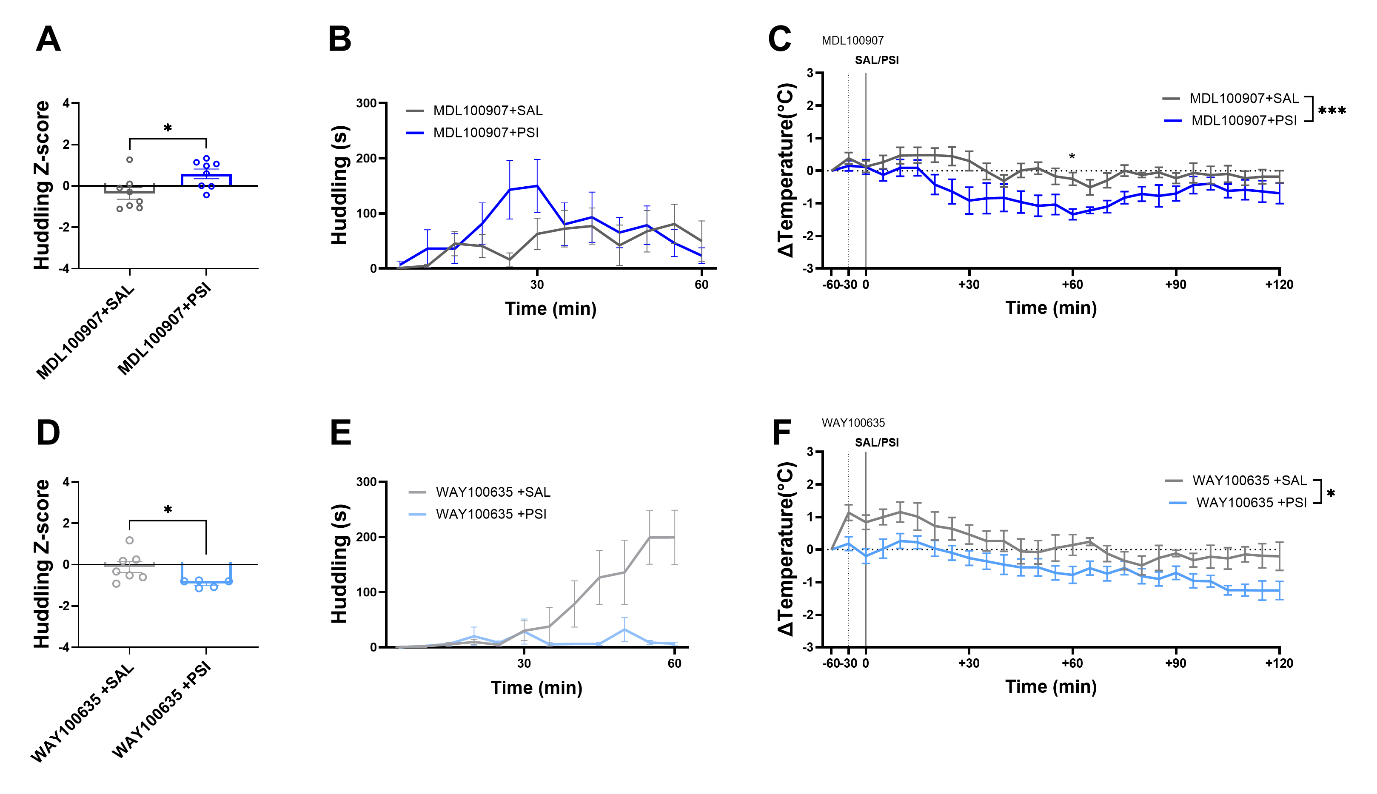

**Supplementary Data 3. Acutely, psilocybin increased huddling in male mice independent of 5-HT2AR antagonism, reduced core body temperature and suppressed huddling when combined with 5-HT1AR antagonism. (A)** PSI increased total huddling behaviour in male mice pre-treated with the 5-HT2AR antagonist (*p* = 0.0194). **(B)** However, no significant differences were observed in huddling behaviour across the 60-minute trial. **(C)** PSI significantly reduced core body temperature in this group, particularly at 60 minutes post-administration (*p* = 0.0365) **(D)** In mice pre-treated with the 5-HT1AR antagonist, PSI significantly reduced total huddling behaviour compared to controls (*p* = 0.0327). **(E)** No significant changes in huddling behaviour were observed across the 60-minute period. **(F)** PSI did not significantly alter core body temperature in 5-HT1AR-antagonised mice. Psilocybin (PSI), saline (SAL), 5-HT2AR antagonist (MDL100907), 5-HT1AR antagonist (WAY100635). Data are presented as mean ± SEM. Statistical analyses were performed using unpaired t-test, mixed-effects models or two-way ANOVA with Šidák post hoc tests. **p*< 0.05.

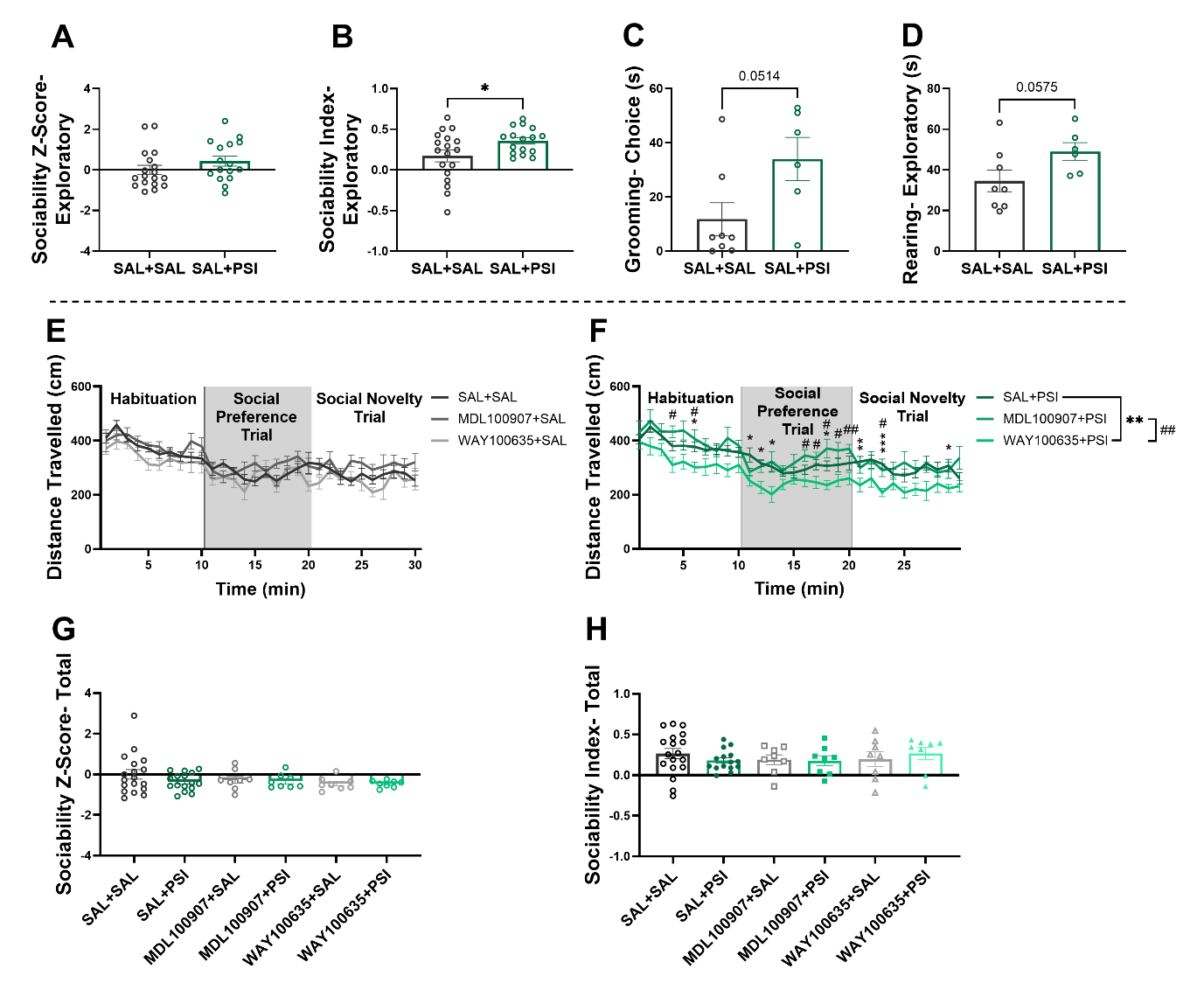

**Supplementary Data 4. Psilocybin modulated sociability and stereotypic behaviour in female mice in a stage-dependent manner at 4 hours post administration during the social novelty trial, while 5-HTR antagonism altered locomotion. (A)** Sociability Z-score was not significantly altered by PSI treatment however, **(B)** PSI-treated mice displayed increased sociability toward the novel mouse compared to the familiar conspecifics (*p =*0.0381). **(C)** These mice also showed a trend toward increased grooming behaviour during the choice phase (*p* = 0.0514), and **(D)** a trend toward increased rearing behaviour during the exploratory phase (*p* = 0.0575). **(E)** 5-HTR antagonism alone did not significantly affect behaviour across the experiment. **(F)** In PSI-treated mice, pre-treatment with 5-HT1AR antagonist significantly reduced locomotor activity compared to the PSI-only group at multiple time points: 6 min (*p* = 0.0227), 11 min (*p* = 0.0188), 12 min (*p* = 0.0123), 13 min (*p* = 0.0256), 18 min (*p* = 0.0236), 21 min (*p* = 0.039), 23 min (*p* = 0.0006), and 29 min (*p* = 0.0165). Compared to the 5-HT2AR antagonist group, the 5-HT1AR antagonist also significantly reduced locomotion at: 4 min (*p* = 0.0107), 6 min (*p* = 0.0465), 16 min (*p* = 0.0297), 17 min (*p* = 0.0278), 18 min (*p* = 0.0151), 19 min (*p* = 0.032), 20 min (*p* = 0.0068), and 23 min (*p* = 0.0329). **(G)** During the social preference trial, neither psilocybin nor 5-HTR antagonism significantly affected the sociability Z-score, **(H)** nor the direct sociability measures. Psilocybin (PSI), saline (SAL), 5-HT2AR antagonist (MDL100907), 5-HT1AR antagonist (WAY100635). Data are presented as mean ± SEM. Statistical analyses were performed using unpaired t-test, one-way ANOVA or two-way ANOVA with Šidák post hoc tests. **p*< 0.05.

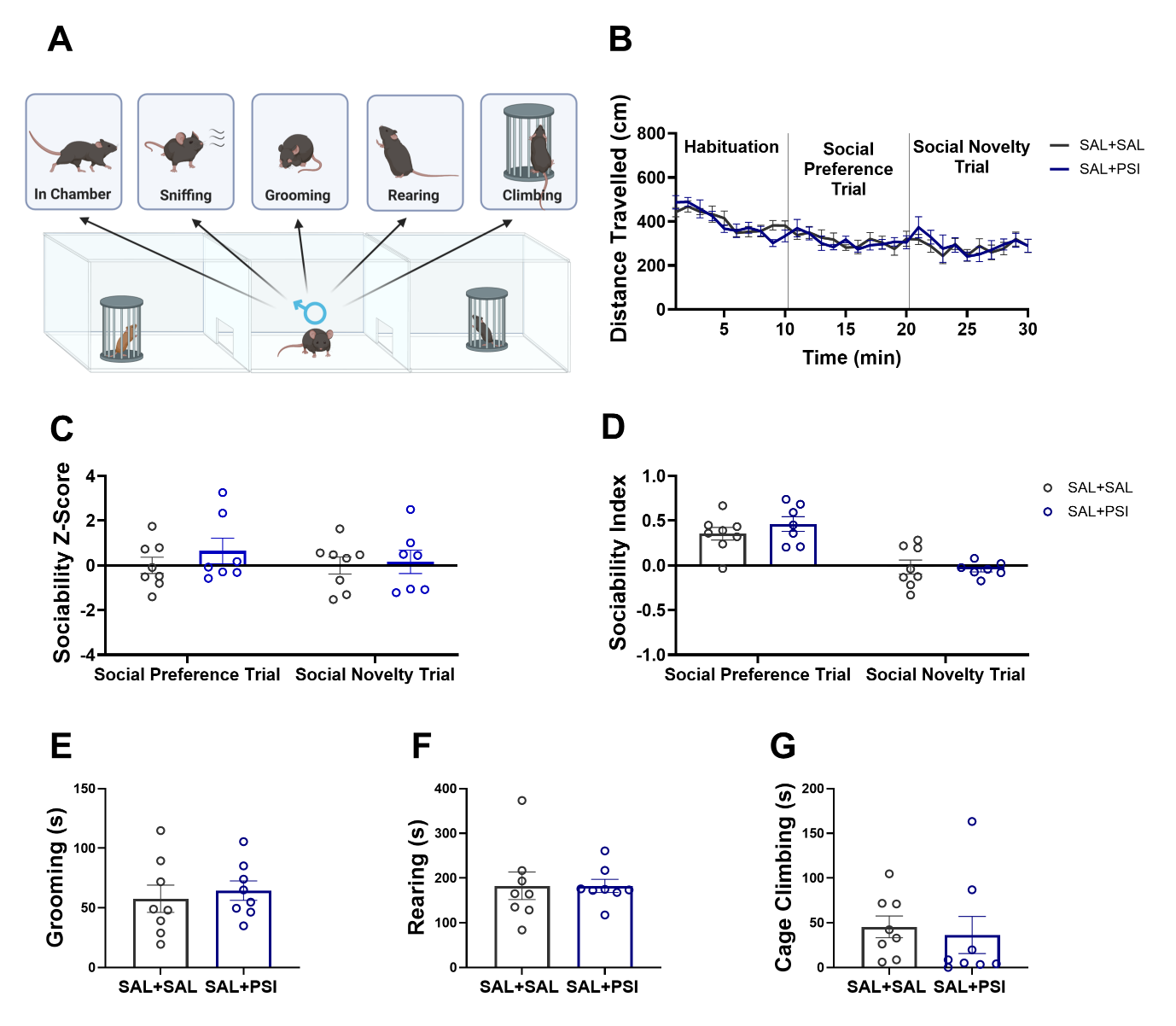

**Supplementary Data 5. Psilocybin did not alter sociability or stereotypic behaviour in male mice 4 hours after administration. (A)** Diagram of the social preference and novelty trial. **(B)** Psilocybin had no effect on locomotor activity, **(C)** sociability Z-score, **(D)** direct sociability, **(E)** grooming, **(F)** rearing or **(G)** cage climbing behaviour in male mice. Psilocybin (PSI), saline (SAL). Data are presented as mean ± SEM. Statistical analyses were performed using unpaired t-test,or two-way ANOVA with Šidák post hoc tests.

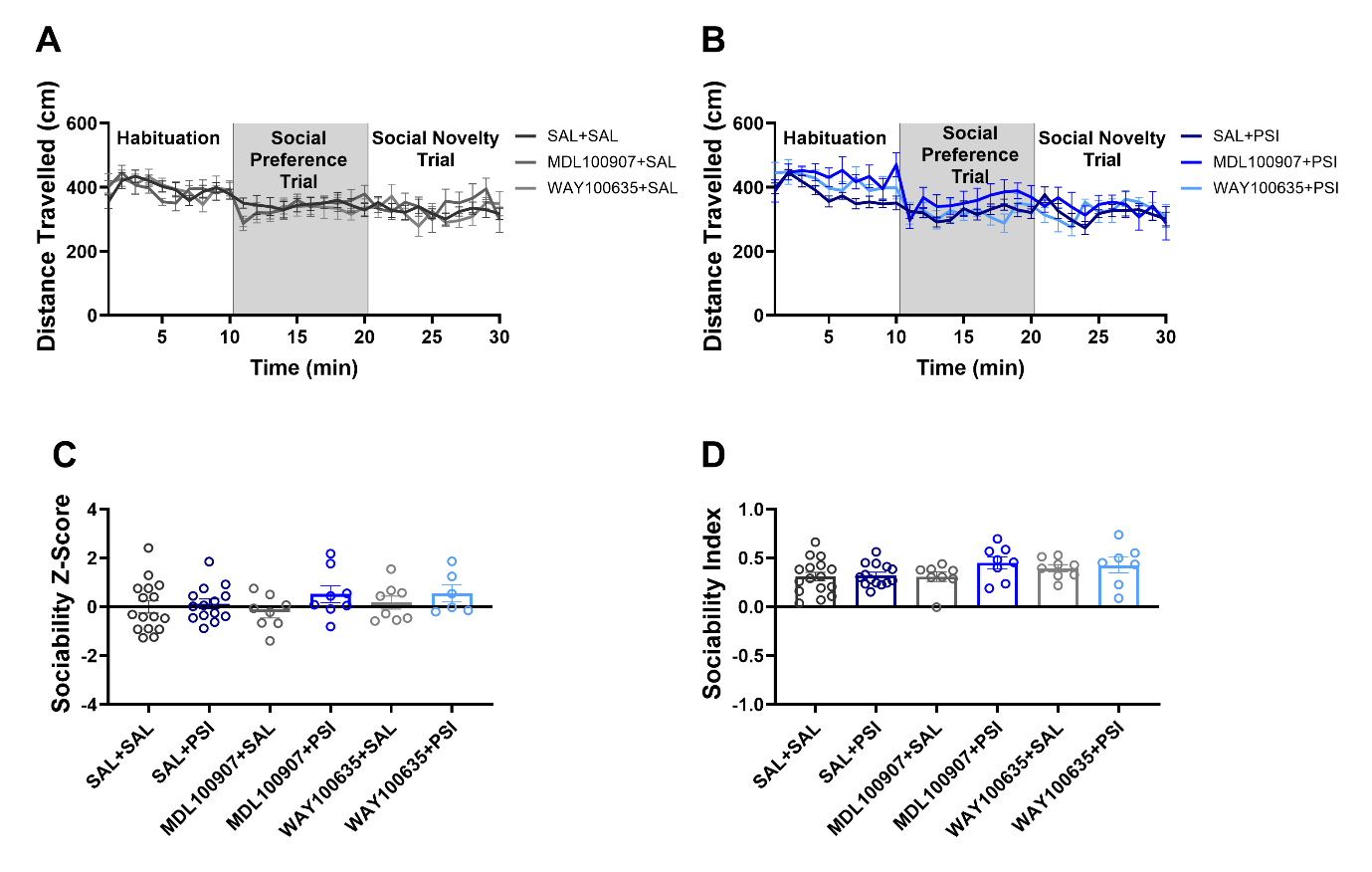

**Supplementary Data 6. Social preference of male mice at 24 hours was not affected by 5-HT receptor antagonism.** 5-HTR antagonism did not alter locomotor activity in SAL-treated male mice, **(B)** nor in PSI-treated males. **(C)** PSI nor 5-HTR antagonism affected the sociability Z-score of male mice during social preference trial, **(D)** and no differences were observed in overall sociability during the same trial. Psilocybin (PSI), saline (SAL), 5-HT2AR antagonist (MDL100907), 5-HT1AR antagonist (WAY100635). Data are presented as mean ± SEM. Statistical analyses were performed using one-way ANOVA or two-way ANOVA with Šidák post hoc tests.

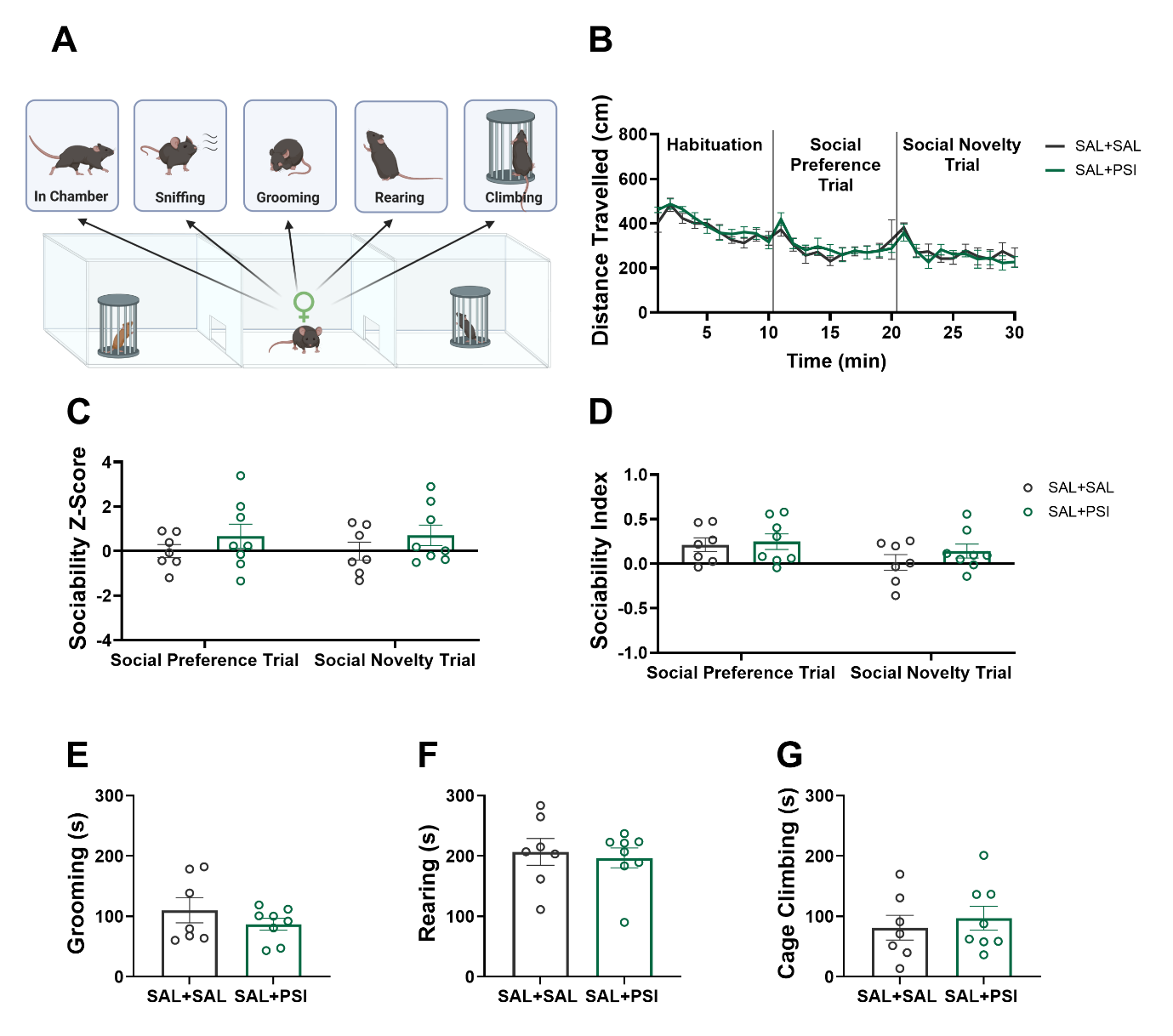

**Supplementary Data 7. Sociability in female mice remained unchanged 24 hours after psilocybin administration. (A)** Schematic of the social preference and novelty trial. **(B)** Psilocybin did not affect locomotor activity, **(C)** sociability Z-score, **(D)** overall sociability, **(E)** grooming, **(F)** rearing or **(G)** cage climbing behaviours in female mice. Psilocybin (PSI), saline (SAL). Data are presented as mean ± SEM. Statistical analyses were performed using unpaired t-test or two-way ANOVA with Šidák post hoc tests.

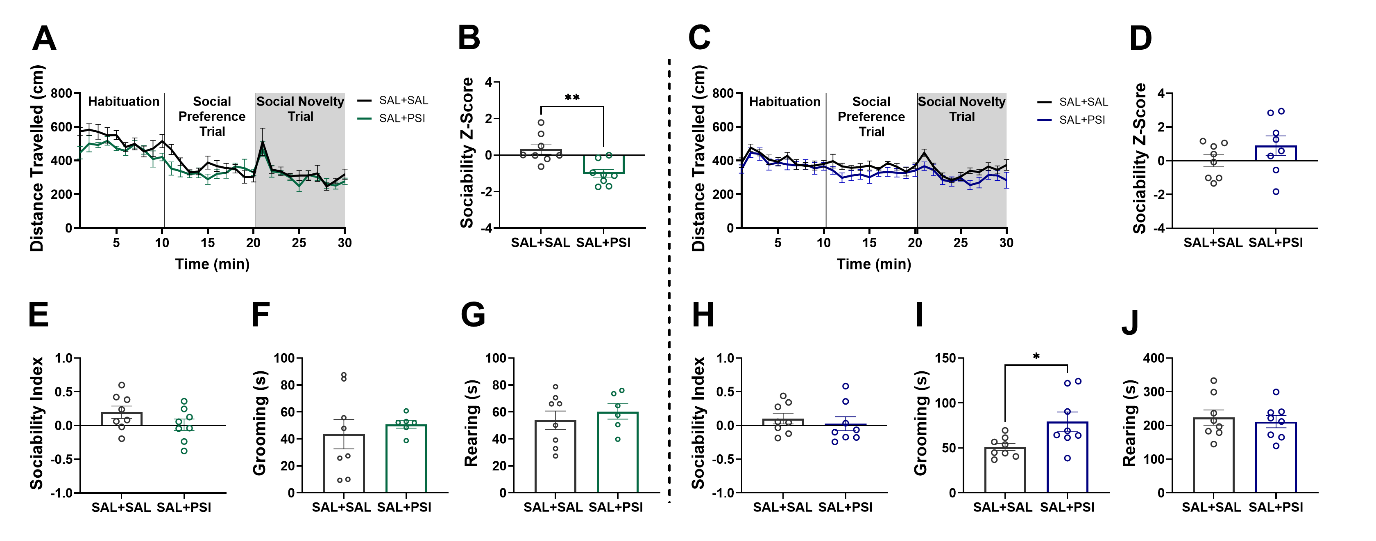

**Supplementary Data 8. Psilocybin increased preference for familiarity in female mice and grooming behaviour in male mice seven days after administration. (A)** Psilocybin did not alter locomotor activity in female mice however, **(B)** it significantly reduced sociability Z-score in these mice (*t* (14) = 3.714, *p* = 0.0023). **(C)** In male mice, psilocybin did not affect locomotor activity and **(D)** or sociability Z-score. **(E)** Psilocybin did not affect sociability index, **(F)** grooming or **(G)** rearing behaviours in female mice. **(H)** Similarly, in male mice psilocybin did not affect the sociability index, **(I)** but significantly increased grooming behaviour (*t* (8.864) = 2.464, *p* = 0.0363) **(J)** without impacting rearing behaviour in these mice. Psilocybin (PSI), saline (SAL). Data are presented as mean ± SEM. Statistical analyses were performed using unpaired t-test or two-way ANOVA with Šidák post hoc tests. **p*< 0.05, ***p*< 0.01.

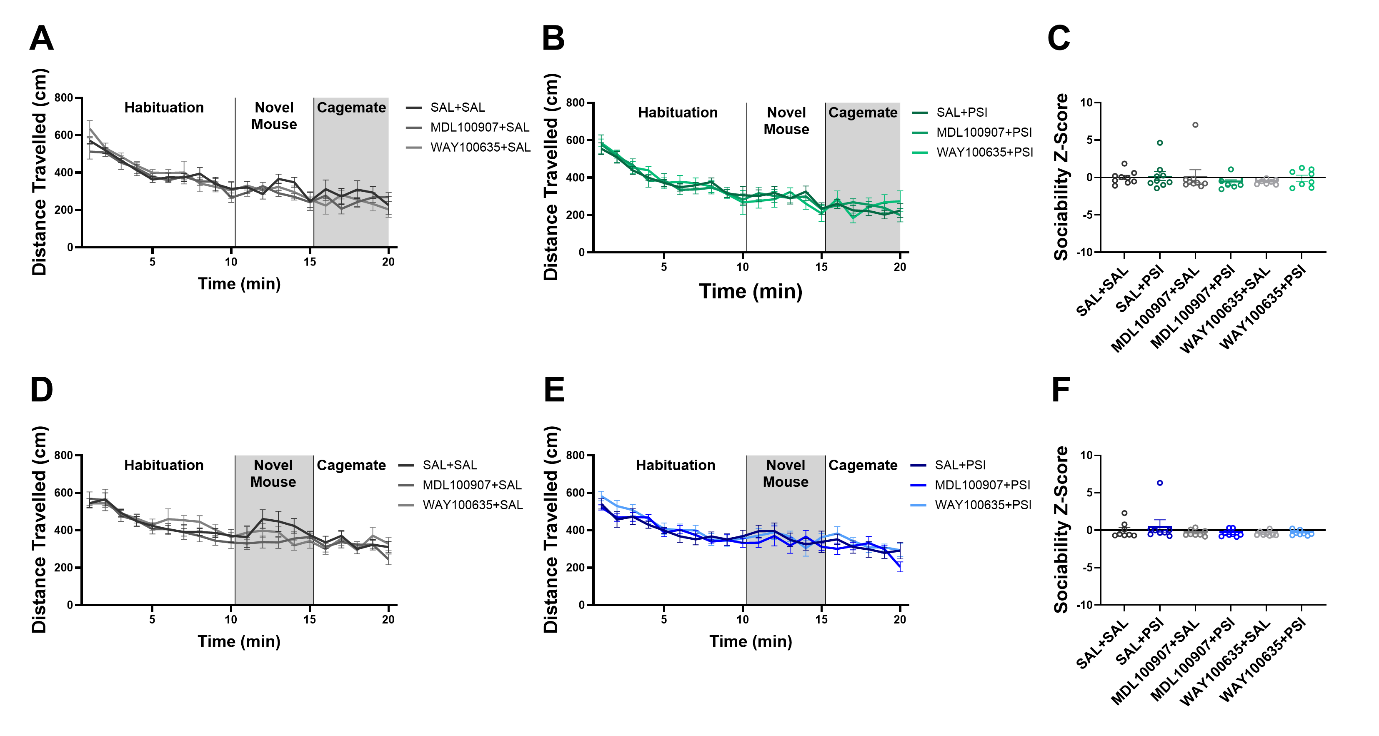

**Supplementary Data 9. Locomotor activity was not affected by 5-HT receptor antagonism during the barrier climbing test at 24 hours, and sociability remained unchanged in a sex-dependent manner. (A)** 5-HTR antagonism did not alter locomotor activity in SAL-treated female mice, **(B)** nor in PSI-treated females. **(C)** Neither PSI nor 5-HTR antagonism affected sociability in female mice. **(D)** Similarly, 5-HTR antagonism did not affect locomotor activity in saline-treated male mice, **(E)** or in the PSI-treated males. **(F)** No effects of PSI or 5-HTR antagonism were observed on sociability in male mice. Psilocybin (PSI), saline (SAL), 5-HT2AR antagonist (MDL100907), 5-HT1AR antagonist (WAY100635). Data are presented as mean ± SEM. Statistical analyses were performed using one-way ANOVA or two-way ANOVA with Šidák post hoc tests.

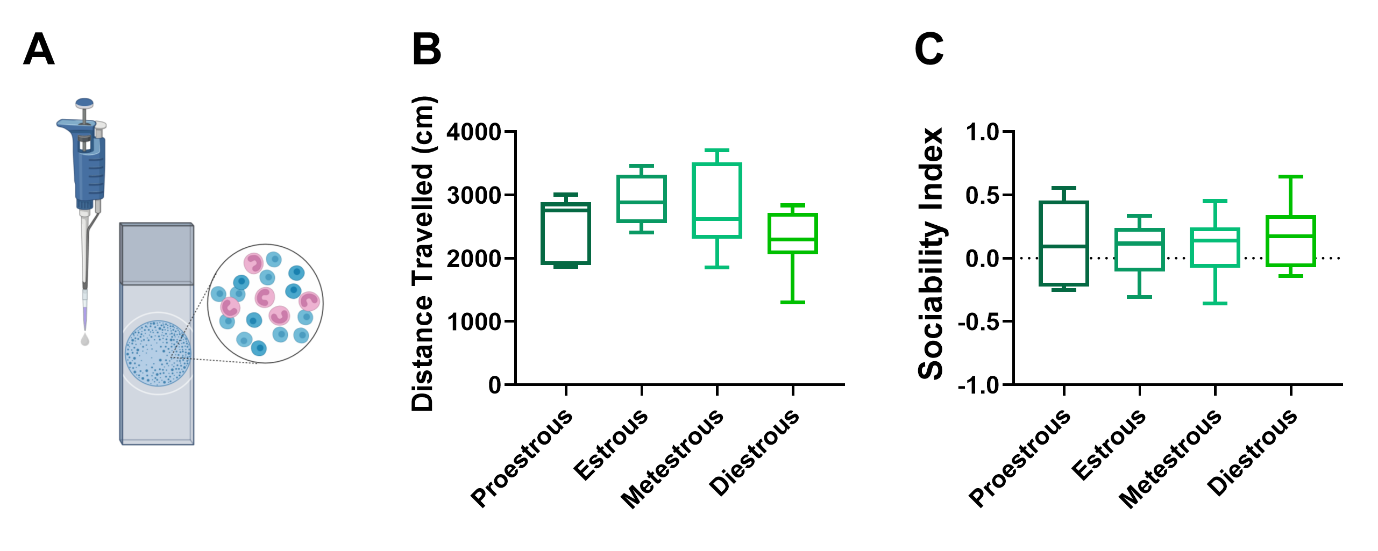

**Supplementary Data 10. No correlation was observed between the stage of the estrous cycle and either locomotor activity or sociability in female mice during the social novelty trial. (A)** Schematic representation of vaginal smear collection, with representative histological images of cell types observed under the microscope. **(B)** The locomotion and **(C)** sociability index did not differ significantly across estrous cycle stages.
Data are presented as mean ± SEM. Statistical analysis was performed using one-way ANOVA.

### **Statistics Table Figure 1.**

| Figure | Statistical test | Group n | Main analysis result | Post-hoc multiple comparisons of interest |
| --- | --- | --- | --- | --- |
| 1.A | t-test | SAL+SAL = 9  SAL+PSI = 9 | t (15.47)= 2.253, p= 0.0391 |  |
| 1.B | Two-way ANOVA | SAL+SAL = 9  SAL+PSI = 9 | Time (F (4.598, 72.32) = 3.136, p= 0.0151); Treatment (F (1, 16) = 7.723, p= 0.0134)  Time x treatment interaction (F (4.598, 72.32) = 3.105, p= 0.0159) | 25 min (= 0.0133), 30 min (p= 0.0155), 45 min (p=0.0311) |
| 1.C | Mixed-effects analysis | SAL+SAL = 9  SAL+PSI = 9 | Time (F (26, 394) = 7.185, p< 0.0001); Treatment (F (1, 16) = 16.1, p= 0.0001)  Time x treatment interaction (F (26, 394) = 5.626, p<0.0001) | 5 min (p= 0.0277), 15 min (p=0.0277), 20 min (p= 0.0004), 25 min (p< 0.0001), 30 min (p< 0.0001), 35 min (p< 0.0001), 40 min (p< 0.0001), 45 min (p< 0.0001), 50 min (p= 0.0001), 55 min (p= 0.0018), 60 min (p= 0.0111), 65 min (p= 0.0026), 70 min (p= 0073) |
| 1.D | One-way ANOVA | SAL+SAL = 9, MDL +SAL = 8, WAY+ SAL = 5, SAL+PSI = 9, MDL+PSI = 8, WAY+ PSI= 8 | All SAL-treated mice: F (2, 19) = 1.684, p= 0.0081  All PSI-treated mice: F (2,21) = 2.098, p< 0.0001 | SAL+SAL vs WAY+SAL: p= 0.0062, SAL+PSI vs MDL+PSI: p= 0.0029,  SAL+PSI vs WAY+ PSI: p< 0.0001 |
| 1.E | Two-way ANOVA | SAL+SAL = 9, MDL +SAL = 8, WAY+ SAL = 5, | Time (F (2.632, 49.53) = 4.398, p= 0.0106) | N/A |
| 1.F | Two-way ANOVA | SAL+PSI = 9, MDL+PSI = 8, WAY+ PSI= 8 | Time (F (4.204, 87.15) = 5.438, p= 0.0005);  Treatment (F (2,21) = 5.604), p= 0.0112) | SAL+PSI vs WAY+PSI: 25 min (p= 0.0228), 30 min (p= 0.0158), 35 min (p= 0.0275), 40 min (p= 0.0428),  SAL+PSI vs MDL+PSI: 50 min (p= 0.0419) |
| 1.G | Mixed-effects analysis | SAL+SAL = 9, MDL +SAL = 8, WAY+ SAL = 5, | Time (F (5.728, 119.4) = 3.327, p= 0.0052), Treatment (F (2.22) = 4.417, p= 0.0244) | SAL+SAL vs WAY+ SAL: 0 min (p= 0.0206), 5 min (p= 0.0398), 25 min (p= 0.0257), 65 min (p= 0.027), 70 min (p= 0.0112)  MDL +SAL vs WAY+ SAL: 45 min (p= 0.0258), 65 min (p= 0.0099) |
| 1.H | Mixed-effects analysis | SAL+PSI = 9, MDL+PSI = 8, WAY+ PSI= 8 | Time (F (3.531, 68.59) = 18.1, p< 0.0001), Time x treatment (F (52, 505) = 1.665, p= 0.0034 | SAL+PSI vs MDL+PSI: 0 min (p= 0.0398), 85 min (p= 0.0429)  SAL+PSI vs WAY+PSI: 75 min (p= 0.021),  MDL+PSI vs WAY+PSI: 75 min (p= 0.0007), 80 min (p= 0.02), 90 min (p= 0.0081), 95 min (p= 0.0319), 110 min (p= 0.025), 115 min (p= 0.0113), 120 min (p= 0.0152) |

| Figure | Statistical test | Group n | Main analysis result | Post-hoc multiple comparisons of interest |
| --- | --- | --- | --- | --- |
| 1.I | t-test | SAL+SAL = 8  SAL+PSI = 9 | N/A | N/A |
| 1.J | Two-way ANOVA | SAL+SAL = 8  SAL+PSI = 9 | Time (F (11,143) = 3.923, p< 0.0001) | N/A |
| 1.K | Mixed-effects analysis | SAL+SAL = 8  SAL+PSI = 9 | Time (F (4.092,56.19) = 10.65, p< 0.0001) | N/A |
| 1.L | One-way ANOVA | SAL+SAL = 8, MDL +SAL = 8, WAY+SAL = 7, SAL+PSI = 9, MDL+PSI = 8, WAY+PSI= 5 | All PSI-treated mice: (F (2,19) = 1.669, p= 0.0007) | SAL+PSI vs WAY+ PSI: p= 0.0018, MDL+PSI vs WAY+ PSI: p= 0.0009 |
| 1.M | Two-way ANOVA | SAL+SAL = 8, MDL+SAL = 8, WAY+SAL = 7 | Time (F (3.625,65.25)= 6.525, p= 0.0003), Time x treatment interaction (F (7.25,65.25)= 2.213, p= 0.0425) | SAL+SAL vs WAY+ SAL: 0 min (p= 0.0399) |
| 1.N | Two-way ANOVA | SAL+PSI = 9, MDL+PSI = 8, WAY+PSI = 5 | Time x treatment interaction (F (7.734,73.47) = 2.186, p= 0.0399 | SAL+PSI vs WAY+ PSI: 45 min (p= 0.0496), 55 min (p= 0.0489) |
| 1.O | Mixed-effects analysis | SAL+SAL = 8, MDL+SAL = 8, WAY+SAL = 7 | Time (F (5.062,96.96) = 16.36, p< 0.0001) | N/A |
| 1.P | Mixed-effects analysis | SAL+PSI = 9, MDL+PSI = 8, WAY+PSI= 5 | Time (F (4.341,81.64) = 13.79, p< 0.0001), Time x treatment interaction (F (52,489) = 1.496, p= 0.0172) | SAL+PSI vs WAY+ PSI: 0 min (p= 0.0289),  SAL+PSI vs MDL+ PSI: 20 min (p= 0.0165) |

### **Statistics Table Figure 2.**

| Figure | Statistical test | Group n | Main analysis result | Post-hoc multiple comparisons of interest |
| --- | --- | --- | --- | --- |
| 2.B | Two-way ANOVA | SAL+SAL = 18  SAL+PSI = 16 | Time (F (11.02,341.9) = 20.85, p< 0.0001) | N/A |
| 2.C | t-test | SAL+SAL = 18  SAL+PSI = 16 | t (32) = 1.71, p= 0.097 | N/A |
| 2.D | t-test | SAL+SAL = 18  SAL+PSI = 16 | t (32) = 2.043, p= 0.0494 | N/A |
| 2.E | t-test | SAL+SAL = 8  SAL+PSI = 6 | t (8.016) = 2.383, p= 0.0443 | N/A |
| 2.F | t-test | SAL+SAL = 8  SAL+PSI = 6 | N/A | N/A |
| 2.G | One-way ANOVA | SAL+SAL = 18, MDL+SAL = 8, WAY+SAL = 8, SAL+PSI = 16, MDL+PSI= 8, WAY+PSI= 8 | All PSI-treated mice: F (2,29) = 1.868, p= 0.0107 | SAL+PSI vs MDL+PSI: p= 0.0252  SAL+PSI vs WAY+PSI: p= 0.023 |
| 2.H | One-way ANOVA | SAL+SAL = 18, MDL+SAL = 8, WAY+SAL = 8, SAL+PSI = 16, MDL+PSI = 8, WAY+PSI= 8 | All PSI-treated mice: F (2,29) = 0.5308, p= 0.0008 | SAL+PSI vs MDL+PSI: p= 0.0007  SAL+PSI vs WAY+PSI: p= 0.0243 |

### **Statistics Table Figure 3.**

| Figure | Statistical test | Group n | Main analysis result | Post-hoc multiple comparisons of interest |
| --- | --- | --- | --- | --- |
| 3.B | Two-way ANOVA | SAL+SAL = 15, SAL+PSI = 16 | Time (F (10.09,282.5) = 8.968, p< 0.0001) | N/A |
| 3.C | t-test | SAL+SAL = 16, SAL+PSI = 14 | N/A | N/A |
| 3.D | t-test | SAL+SAL = 16, SAL+PSI = 14 | N/A | N/A |
| 3.E | t-test | SAL+SAL = 7, SAL+PSI = 8 | t (7.119) = 3.145, p= 0.0159 | N/A |
| 3.F | t-test | SAL+SAL = 7, SAL+PSI = 8 | t (10.8) = 3.744, p= 0.0033 | N/A |
| 3.G | One-way ANOVA | SAL+SAL = 16, MDL+SAL = 8, WAY+SAL = 8, SAL+PSI = 14, MDL+PSI = 7, WAY+PSI= 7 | All SAL-treated mice: F (2,29) = 0.3241, p= 0.0287 | SAL+SAL vs MDL +SAL: p= 0.0168 |
| 3.H | One-way ANOVA | SAL+SAL = 16, MDL+SAL = 8, WAY+SAL = 8, SAL+PSI = 14, MDL+PSI = 7, WAY+PSI= 7 | All SAL-treated mice: F (2,29) = 2.159, p= 0.0141  SAL+PSI vs WAY+ PSI: t (11.17) = 2.108, p= 0.0584 | SAL+SAL vs MDL +SAL: p= 0.0077 |

### **Statistics Table Figure 4.**

| Figure | Statistical test | Group n | Main analysis result | Post-hoc multiple comparisons of interest |
| --- | --- | --- | --- | --- |
| 4.A | Two-way ANOVA | SAL+SAL = 10, SAL+PSI = 9 | Time (F (6.184,102.8) = 20.08, p< 0.0001) | N/A |
| 4.B | Two-way ANOVA | SAL+SAL = 10, SAL+PSI = 8 | N/A | N/A |
| 4.C | t-test | SAL+SAL = 10, SAL+PSI = 9 | t (17) = 2.045, p= 0.0566 | N/A |
| 4.D | One-way ANOVA | SAL+SAL = 10, MDL+SAL = 9, WAY+SAL = 8, SAL+PSI = 8, MDL+PSI = 8, WAY+PSI= 7 | N/A | N/A |
| 4.E | One-way ANOVA | SAL+SAL = 10, MDL+SAL = 9, WAY+SAL = 7, SAL+PSI = 8, MDL+PSI= 8, WAY+PSI= 8 | All PSI-treated mice: F (5,44) = 6.315, p= 0.0108 | SAL+PSI vs MDL+PSI: p= 0.0694 |
| 4.F | Two-way ANOVA | SAL+SAL = 8, SAL+PSI = 8 | Time (F (6.424,89.6) = 10.81, p< 0.0001) | N/A |
| 4.G | Two-way ANOVA | SAL+SAL = 8, SAL+PSI = 8 | Phase (F (2,42) = 18.77, p< 0.0001) | Habituation (p= 0.0407) |
| 4.H | t-test | SAL+SAL = 8, SAL+PSI = 8 | t (14) = 2.335, p= 0.0349 | N/A |
| 4.I | One-way ANOVA | SAL+SAL = 8, MDL+SAL = 8, WAY+SAL = 8, SAL+PSI = 8, MDL+PSI = 7, WAY+PSI= 7 | N/A | N/A |
| 4.J | One-way ANOVA | SAL+SAL = 8, MDL+SAL = 8, WAY+SAL = 8, SAL+PSI = 8, MDL+PSI = 7, WAY+PSI= 7 | N/A | N/A |

### **Statistics Table Figure 5.**

| Figure | Statistical test | Group n | Main analysis result | Post-hoc multiple comparisons of interest |
| --- | --- | --- | --- | --- |
| 5.C | Mixed-effects analysis  Inset: Two-way ANOVA | SAL = 5, PSI= 5 | Familiar Mouse: N/A  Novel Mouse:  Inset: | N/A |
| 5.D | Two-way ANOVA | SAL = 5, PSI= 5 | N/A | N/A |
| 5.E | Mixed-effects analysis  Inset: Two-way ANOVA | SAL = 4, PSI= 5 | Familiar Mouse: Treatment (F (1,7) = 9.056, p= 0.0197), Treatment x Mouse (F (3.647,25.43) = 3.221, p= 0.032)  Inset: F (1,14) = 9.169, p= 0.009 | Inset: Familiar Mouse: p= 0.0317 |
| 5.F | Two-way ANOVA | SAL = 4, PSI= 5 | N/A | N/A |
| 5.G | Two-way ANOVA | SAL = 4, PSI= 3 | N/A | N/A |
| 5.H | Two-way ANOVA | SAL = 4, PSI= 3 | Peak Z-Score: Treatment (F (1,10) = 4.994, p= 0.0494) | Novel Mouse: SAL vs PSI: p= 0.086 |
| 5.I | Mixed-effects analysis  Inset: Two-way ANOVA | SAL = 4, PSI= 3 | N/A | N/A |
| 5.J | Two-way ANOVA | SAL = 4, PSI= 3 | Mean Z-Score: Treatment (F (1,14) = 4.059, p= 0.0636),  Peak Z-Score: Treatment x Mouse interaction (F (1,14) = 5.131, p= 0.399) | Novel Mouse: SAL vs PSI: p= 0.0326  SAL: Familiar vs Novel Mouse: p= 0.0353 |

### **Statistics Table- Supplementary Data 1.**

| Figure | Statistical test | Group n | Main analysis result | Post-hoc multiple comparisons of interest |
| --- | --- | --- | --- | --- |
| S1.A | One-way ANOVA | SAL+SAL = 9, MDL+SAL = 8, WAY+SAL = 5, SAL+PSI = 9, MDL+PSI = 8, WAY+PSI = 7 | All SAL-treated mice: F (2,19) = 1.19, p= 0.0364  All PSI-treated mice: F (2,21) = 0.6772, p= 0.0366 | SAL+SAL vs MDL +SAL: p= 0.0339  SAL+PSI vs MDL+PSI: p= 0.0328 |
| S1.B | One-way ANOVA | SAL+SAL = 8, MDL+SAL = 8, WAY+SAL = 5, SAL+PSI = 9, MDL+PSI = 8, WAY+PSI = 7 | N/A | N/A |
| S1.C | One-way ANOVA | SAL+SAL = 8, MDL+SAL = 8, WAY+SAL = 7, SAL+PSI = 9, MDL+PSI = 8, WAY+PSI = 5 | N/A | N/A |
| S1.D | One-way ANOVA | SAL+SAL = 8, MDL+SAL = 8, WAY+SAL = 7, SAL+PSI = 6, MDL+PSI = 8, WAY+PSI = 4 | All SAL-treated mice: F (2,19) = 1.343, p= 0.0069 | SAL+SAL vs MDL +SAL: p=0.0366,  SAL+SAL vs WAY+ PSI: p= 0.0101 |

### **Statistics Table- Supplementary Data 2.**

| Figure | Statistical test | Group n | Main analysis result | Post-hoc multiple comparisons of interest |
| --- | --- | --- | --- | --- |
| S2.A | t-test | MDL+SAL= 8, MDL+PSI = 8 | N/A | N/A |
| S2.B | Two-way ANOVA | MDL+SAL= 8, MDL+PSI = 8 | Time (F (3.502,48.39) = 3.448, p= 0.0186), Time x treatment interaction (F (3.502,48.39) = 4.423, p< 0.0056) | N/A |
| S2.C | Mixed-effects analysis | MDL+SAL= 8, MDL+PSI = 8 | Time (F (3.002,36.71) = 6.813, p= 0.0009), Treatment (F (1.14) = 8.965, p= 0.0097), Time x treatment interaction (F (3.002,36.71) = 4.398, p= 0.0096) | 45 min (p= 0.0297) |
| S2.D | t-test | MDL+SAL = 5, MDL+PSI= 7 | N/A | N/A |
| S2.E | Two-way ANOVA | WAY+SAL= 5, WAY+PSI = 7 | N/A | N/A |
| S2.F | Mixed-effects analysis | WAY+SAL= 8, WAY+PSI = 8 | Time (F (2.577,33.2) = 6.192, p= 0.0028), Treatment (F (1.14) = 5.376, p= 0.0361), Time x treatment interaction (F (26,355) = 3.114, p< 0.0001) | 75 min (p= 0.0446), 90 min (p= 0.0478) |

### **Statistics Table- Supplementary Data 3.**

| Figure | Statistical test | Group n | Main analysis result | Post-hoc multiple comparisons of interest |
| --- | --- | --- | --- | --- |
| S3.A | t-test | MDL+SAL= 8, MDL+PSI = 8 | t (14) = 2.581, p= 0.0218 | N/A |
| S3.B | Two-way ANOVA | MDL+SAL= 8, MDL+PSI = 8 | Time (F (11,154) = 2.159, p= 0.0194) | N/A |
| S3.C | Mixed-effects analysis | MDL+SAL= 8, MDL+PSI = 8 | Time (F (3.01,36.12) = 5.715, p= 0.0026), Treatment (F (1,14) = 8.987, p= 0.0096), Time x treatment (F (26,312) = 1.602, p= 0.0342) | 60 min (p= 0.0365) |
| S3.D | t-test | MDL+SAL= 7, MDL+PSI = 5 | t (10) = 2.478, p= 0.0327 | N/A |
| S3.E | Two-way ANOVA | WAY+SAL = 5, WAY+PSI = 7 | Time (F (2.585,25.85) = 4.325, p= 0.0169), Treatment (F (1,10) = 7.749, p= 0.0193), Time x treatment (F (2.585,25.85) = 4.067, p= 0.0211) | N/A |
| S3.F | Mixed-effects analysis | WAY+SAL= 8, WAY+PSI = 8 | Time (F (3.559,43.53) = 18.03, p< 0.0001), Treatment (F (1,14) = 6.696, p= 0.0215) | N/A |

### **Statistics Table- Supplementary Data 4.**

| Figure | Statistical test | Group n | Main analysis result | Post-hoc multiple comparisons of interest |
| --- | --- | --- | --- | --- |
| S4.A | t-test | SAL+SAL = 18  SAL+PSI = 16 | N/A | N/A |
| S4.B | t-test | SAL+SAL = 18  SAL+PSI = 16 | t (32) = 2.164, p= 0.0381 | N/A |
| S4.C | t-test | SAL+SAL = 8  SAL+PSI = 6 | t (10.14) = 2.208, p= 0.0514 | N/A |
| S4.D | t-test | SAL+SAL = 8  SAL+PSI = 6 | t (11.99) = 2.1, p= 0.0575 | N/A |
| S4.E | Mixed-effects analysis | SAL+SAL = 18, MDL+SAL = 8, WAY+ SAL = 8 | Time (F (11.85,348.9) = 17.17, p< 0.0001) | N/A |
| S4.F | Mixed-effects analysis | SAL+PSI = 16, MDL+PSI = 8, WAY+PSI = 8 | Time (F (10.05,286.9) = 18.04, p< 0.0001), Treatment (F (2.29) = 6.626, p= 0.0043) | MDL+PSI vs WAY+ PSI: 4 min (p= 0.0107), 6 min (p= 0.0465), 16 min (p= 0.0297), 17 min (p= 0.0278), 18 min (p= 0.0151), 19 min (p= 0.032), 20 min (p= 0.0068), 23 min (p= 0.0329)  SAL+PSI vs WAY+ PSI: 6 min (p= 0.0227), 11 min (p= 0.0188), 12 min (p= 0.0123), 13 min (p= 0.0256), 18 min (p= 0.0236), 21 min (p= 0.039), 23 min (p= 0.0006), 29 min (p= 0.0165) |
| S4.G | One-way ANOVA | SAL+SAL = 18, MDL+SAL = 8, WAY+ SAL = 8, SAL+PSI = 16, MDL+PSI = 8, WAY+PSI = 8 | F (5,60) = 3.001, p= 0.0176 | N/A |
| S4.H | One-way ANOVA | SAL+SAL = 18, MDL+SAL = 8, WAY+SAL = 8, SAL+PSI = 16, MDL+PSI = 8, WAY+PSI = 8 | N/A | N/A |
